# *In vivo* models for rustrela virus (RusV) infection in potential reservoir and spill-over hosts

**DOI:** 10.1101/2025.11.10.687696

**Authors:** Antonia Klein, Lukas Mathias Michaely, Jessica Geers, Sophie-Celine Weinert, Claudia Wylezich, Andrea Aebischer, Lorenz Ulrich, Martin Beer, Kore Schlottau, Angele Breithaupt, Dennis Rubbenstroth

**Affiliations:** Institute of Diagnostic Virology, Friedrich-Loeffler-Institut, Greifswald-Insel Riems, Germany; Department of Experimental Animal Facilities and Biorisk Management, Friedrich-Loeffler-Institut, Greifswald-Insel Riems, Germany

**Author notes:** Corresponding author: Dennis Rubbenstroth.

**Keywords:** rustrela virus, RusV, experimental infection, animal model, wood mouse, rat, lymphohistiocytic encephalitis, reproduction of disease

## Abstract

Rustrela virus (RusV; species *Rubivirus strelense*) is a recently discovered relative of the human rubella virus (*Rubivirus rubellae*) and causes fatal non-suppurative meningoencephalomyelitis in a broad range of mammalian domestic, wild and zoo animals, in Europe and the USA, including ‘staggering disease’ in domestic cats. Based on its reportedly broad host range, a zoonotic potential of RusV cannot be excluded. Apparently healthy yellow-necked field mice (*Apodemus flavicollis*) and wood mice (*Apodemus sylvaticus*) were identified as potential reservoir hosts.

In this study, we experimentally inoculated wood mice and Lewis rats with brain homogenates of RusV-infected animals. Intracranial as well as combined intranasal/peroral inoculation resulted in apparently persistent RusV infection in both rodent species, while combined intramuscular/subcutaneous injection failed to establish detectable infection. Viral loads were highest in the central nervous system, but viral RNA was prominently found also in mucosal organs of wood mice. Viral RNA was frequently detected in oral swabs from infected wood mice, but only sparsely from rats. Neither wood mice nor rats presented neurological signs during twelve weeks after infection. However, RusV-infected rats showed significantly reduced body weight. A lymphohistiocytic meningoencephalomyelitis was observed in infected rats, whereas infected wood mice infrequently developed minimal lesions in the brain.

Our study provides the first infection model for RusV and indicates that natural infection may occur via mucosal routes. Shedding of viral RNA suggests wood mice to serve as natural reservoir hosts. In rats, we could reproduce a lymphohistiocytic meningoencephalomyelitis, as described for diseased cats and other tentative non-reservoir hosts.

## INTRODUCTION

Rustrela virus (RusV; species *Rubivirus strelense*; family *Matonaviridae*) is a relative of rubella virus (RuV; *Rubivirus rubella*e), the causative agent of German measles in humans. In 2020, RusV was discovered by metagenomic sequencing of samples from zoo animals of various species suffering from lymphohistiocytic encephalitis in a zoological garden in northeastern Germany as one of the first known relatives of RuV [1–4]. Subsequent surveys of fresh or archived brain samples from animals with encephalitis of unknown cause confirmed the presence of different genetic RusV variants also in zoo animals and domestic cats in other parts of Germany [4–7], in domestic cats in Sweden and Austria [5, 8, 9] and even in a mountain lion in Colorado, USA [10]. These studies connected RusV also to two long-known disease entities, “staggering disease” and “lion encephalitis”, the cause of which had remained obscure for more than five decades [5–7]. Staggering disease was first described in the 1970s in domestic cats in Sweden and is still incident in a hotspot area around Lake Malaren [5, 9, 11]. Furthermore, it had been described in an area close to Vienna in Austria in the early 1990s but appears to have vanished since [5, 8, 12, 13]. Lion encephalitis had been reported in zoos in Germany mainly in the 1970s and 1980s [6, 7, 14, 15]. Previous assumptions that staggering disease was caused by Borna disease virus 1 (BoDV-1) could not be confirmed [5, 8, 9]. Instead, the consistent detection of RusV in the brains of encephalitic animals by different independent methods, in combination with the absence of RusV in non-encephalitic controls strongly indicated RusV as the causative agent of these diseases [5–7, 9]. However, reproduction of the disease to fulfil Henle-Koch’s postulates has not been reported, yet.

In the encephalitic animals, RusV is highly neurotropic. Detection of viral RNA by semiquantitative reverse transcription polymerase chain reaction (RT-qPCR) can be observed predominantly in the central nervous system (CNS), while only very low levels are inconsistently detected in peripheral organs [1, 4, 6, 9]. Infected cells, as identified by RNA *in situ* hybridization (RNA-ISH) and immunohistochemistry (IHC), are mainly neurons and occasionally astrocytes and microglial cells [1, 3–6, 9]. Microscopic lesions reveal a usually multifocal lymphohistiocytic meningoencephalomyelitis of varying intensity with a focus on the grey matter. Inflammatory lesions are characterized by mononuclear perivascular cuffs identified as mainly lymphocytes as well as microglial and astrocyte activation. Notably, these lesions do usually only partly overlap with the areas of the most prominent detection of viral RNA and antigen [1, 3–7, 9, 10]. RusV-associated disease is best described for cats suffering from staggering disease. Hindleg weakness and ataxia leading to a staggering gait are the most consistent clinical signs. Further signs include fever, behavioural changes, hyperaesthesia, reduced reflexes, the inability to retract the claws and seizures [5, 10]. While zoo animals are frequently reported to have died from the disease, domestic cats are usually euthanized due to the progressing or at least persisting neurological disorders and the poor prognosis at few days to several months after disease onset [1, 3–10].

The tissue distribution pattern of RusV suggests that encephalitic cats and zoo animals are unable to shed the virus and act as dead-end hosts [1, 4, 5]. This assumption is further substantiated by phylogeographic analyses demonstrating RusV to be divided into phylogenetic clades and genotypes, which are regionally associated with only partly overlapping dispersal areas [5–7, 16, 17]. Such pattern is typical for pathogens being bound to a rather immobile reservoir host, such as small mammals, while it is incompatible with being spread by much more mobile and regularly traded hosts, such as domestic and zoo animals. Screening of rodents and other small mammals identified RusV to be widely distributed in yellow-necked field mice (*Apodemus flavicollis*) in Northern Germany, while it was not detected in any other tested species [1, 3, 16]. In contrast, in Sweden RusV was found only in wood mice (*Apodemus sylvaticus*; [5]). Inflammatory lesions were not observed in the brains of these animals, suggesting that they might serve as a healthy reservoir host. However, their RusV tissue distribution was similar to those of the supposed dead-end hosts, with viral RNA predominantly detected in the brain.Thus, their role as reservoir as well as their ability to shed and transmit the virus still needs further confirmation [1, 5, 16].

The aim of this study was to establish first *in vivo* infection models for RusV to characterize routes and course of infection, viral shedding and the development of microscopic lesions and possibly clinical disease. Wood mice were selected as a potential reservoir host, whereas rats represented a possible non-reservoir host. Since virus isolation in cell culture has not been successful for RusV, so far, the experimental infection was performed by using brain homogenates originating from RusV-infected animals.

## MATERIAL & METHODS

### Experimental design and animals

In three consecutive experiments (Exp. 1 to 3), subadult wood mice and/or juvenile Lewis rats were inoculated via different routes with brain homogenates originating from naturally or experimentally RusV-infected animals. At weekly intervals, body weights were determined and oral swabs were collected from all animals. At the same time points, pooled faecal samples were collected from each cage of four to five animals that had received the same inoculum. In Exp. 3, environmental swabs were in addition taken from the walls of each cage. Animals were scheduled to be euthanized after four to twelve weeks post inoculation (p.i.). Serum samples were collected from all animals during euthanasia. At necropsy, tissue samples were collected and stored frozen at -80°C until further processing for RNA extraction. In Exp. 2 and 3, additional tissue samples were formalin-fixed for histopathological analysis.

In Exp. 1, seven separately housed groups (A to G) of four wood mice each were inoculated intracranially (i.c.) with brain homogenates originating from naturally RusV-infected animals. The inocula represented putative reservoir hosts (yellow-necked field mice and wood mice) as well as encephalitic spill-over hosts (zoo animals and domestic cats), different countries of origin (Germany, Slovakia, Sweden and Austria) and different RusV genotypes following the nomenclature established by Pfaff et al. [17] (Supplemental Table S1). The experiment was terminated at 4 weeks p.i.

In Exp. 2, five separately housed groups of four Lewis rats each were i.c.-inoculated either with the same brain homogenates from naturally RusV-infected animals that had already been used for the inoculation of groups D and E in Exp. 1 (groups D1 and E1, respectively) or with pooled brain homogenates originating from the RusV-positive wood mice of groups B, D and E (groups B2, D2 and E2, respectively; Supplemental Table S2) of Exp. 1. A sixth group of two animals remained uninfected. All rats were euthanized at 4 weeks p.i.

In Exp. 3, wood mice and Lewis rats were inoculated via different routes with pooled brain homogenates of all RusV-positive animals of groups D of Exp. 1 and D1 and D2 of Exp. 2, which had been inoculated with virus originating from the infected cat NRL.22_007-6 from Sweden [5]. Wood mice and rats were divided into three groups of 15 animals. One group of each species was inoculated by i.c. route, one group each received the virus by combined intramuscular and subcutaneous (i.m./s.c.) injection and one group each received a combined intranasal and peroral (i.n./p.o.) application of the inoculum. From each group, a subgroup of five animals was euthanized at 4, 8 and 12 weeks p.i. A fourth group of each species, consisting of 10 animals each, was mock-inoculated and received brain homogenates from non-inoculated rats from Exp. 2 (Supplemental Table S3).

Inocula were prepared by homogenization of 10 % brain tissue in sterile phosphate-buffered saline (PBS) and subsequently centrifuged for 10 minutes at 10,000 x *g*. Supernatants were collected and stored at -80 °C until used. Anaesthesia for i.c. inoculation was induced by a combination of ketamine and xylazine (50 mg/kg and 10 mg/kg, respectively, intraperitoneally) and the animals were treated orally with meloxicam after the inoculation. All other inoculation procedures were performed under short-term isoflurane anaesthesia. For i.c. inoculation, 10 µl inoculum per animal were injected sagittally in the direction of the prefrontal cortex. I.m./s.c.-inoculated animals received 10 µl inoculum each into the right knee fold and the left femoral muscle. Mucosal inoculation was achieved by pipetting 10 µl of inoculum into the nostrils (i.n.) and in addition into the mouth (p.o.).

All brain tissues used for inoculation in Exp. 1 were analysed by metagenomic sequencing as described by Pfaff et al. [3]. To test for the presence of any microorganisms other than RusV, the sequencing datasets were analysed using the RIEMS tool for taxonomic read assignment [18]. Analysis of 10^6.1^ to 10^6.8^ sequence reads per sample did not indicate the presence of any potential pathogen other than RusV, with the exception of sample BRA-220 with 10^4.6^ reads classified as *Staphylococcus aureus* (data not shown).

Juvenile Lewis rats and subadult wood mice were purchased from commercial breeders and kept at the animal facility of the Friedrich-Loeffler-Institut at biosecurity level (BSL) 2. Holding and experimental conditions followed the European guidelines on animal welfare and care according to the authority of the Federation of European Laboratory Animal Science Associations (FELASA) and the experiments were approved by the federal animal welfare authorities (reference no. 7221.3-2-010/18, 7221.3-2-010/23, 7221.3-1-017/24).

### RNA extraction and RusV-specific RT-qPCR

For RNA extraction, tissue samples and pooled faeces were homogenized in PBS, whereas oral swabs from Exp. 1 and 2 were vortexed in 500 µl PBS. After centrifugation, 100 µl of the supernatant were transferred to a 96 well multititer plate and RNA extraction was performed using the Nucleo Mag Vet Kit (Macherey & Nagel, Düren, Germany) with the KingFisher™ Flex Purification System (Thermo Fisher Scientific, Darmstadt, Germany) according to the manufactureŕs instructions. A modified protocol was used for RNA extraction from swabs collected in Exp. 3. Swabs were vortexed in 500 µl AVL lysis buffer with proteinase K (Qiagen), incubated at 37°C for 30 minutes and briefly centrifuged before 220 µl of the supernatant were transferred to the multititer plate for RNA extraction.

RusV-specific RNA was detected by RT-qPCR panRusV-2a, as described previously [9]. Briefly, RT-qPCR was performed with AgPath-ID One-Step RT-PCR reagents (Thermo Fisher Scientific), primers RusV_234+ (5’-CCCCGTGTTCCTAGGCAC-3’) and RusV_323- (5’-TCGCCCCATTCWACCCAATT-3’; final concentration: 0.8 μM each) and probe RusV_257_P ([FAM]-5’-TGAGCGACCACCCAGCACTCCA-3’-[BHQ1]; 0.4 μM), eGFP primers (0.2 μM each) and probe (0.15 μM) [19], and 2.5 μl extracted RNA in a total volume of 12.5 μl. The reaction was performed with the following cycler setup: 45 °C for 10 min, 95 °C for 10 min, 45 cycles of 95 °C for 15 s, 60 °C for 30 s and 72 °C for 30 s. A standard RNA preparation of a RusV-positive donkey brain [1] served as positive control and was used for the calibration of Cq values in each RT-qPCR analysis.

### Detection of RusV-reactive antibodies by immunofluorescence antibody test (IFAT)

Baby hamster kidney (BHK) cells were seeded into 96 well microtiter plates and simultaneously transfected with pEXPR-IBA103 plasmids (IBA Lifesciences, Göttingen, Germany) containing codon-optimized sequences encoding for the capsid protein [5] or envelope glycoproteins E2/E1 based on the RusV sequence detected in the brain of donkey NRL.19_041-1 (MN552442.2) from Northern Germany [1]. The transfection was performed using JetPEI following the manufactureŕs instructions. After 24 hours, confluent layers of transfected and non-transfected BHK cells were fixed with 3 % formaldehyde and permeabilized with 50 µl of 0.5 % Triton X-100 in PBS for 10 minutes each. Protein expression in the transfected cells was confirmed by immunofluorescence staining using monospecific polyclonal rabbit hyperimmunesera directed against the RusV capsid or E2/E1 proteins (internal IDs #19/20 and #21, respectively).

For endpoint titration of RusV-reactive antibodies in rat and wood mouse sera, two-fold dilution series of the samples were prepared in Tris-HCl buffer with Tween 20 (T9039; Sigma-Aldrich, Schnelldorf, Germany), starting with a 20-fold dilution. 50 µl of each dilution were added in parallel to the permeabilized RusV-positive and - negative wells. After incubation for one hour, the plates were washed three times with PBS, followed by incubation with goat-anti-mouse-IgG Cy3 or goat-anti-rat-IgG Cy3 conjugate (Jackson Immunoresearch, Ely, UK) for another hour. After a final washing step, the assays were analysed by fluorescence microscopy. For each serum dilution, RusV-positive and -negative wells were compared. Wells were considered positive, if approximately 10-20 % antigen-expressing cells in the transfected wells stained markedly brighter than the background staining of non-transfected cells. Endpoint titres were calculated as the reciprocal dilution factor of the highest dilution providing a positive signal. The mouse monoclonal anti-capsid antibody 2H11B1 [5] served as the positive control for the IFAT staining.

In a first screening step, serum samples from all animals were screened using cells co-transfected with RusV capsid- and E2/E1-encoding plasmids. In a second step, all sera that had tested positive in the initial screening were tested using cells individually transfected with either capsid- or E2/E1-encoding plasmids.

### Histopathology

A tissue sample spectrum including nose, brain, spinal cord, eye, adrenal gland, lung, heart, kidney, spleen, liver, mesenteric lymph nodes, salivary gland, pancreas, stomach, small intestine, large intestine and urinary bladder of wood mice (Exp. 3) and rats (Exp. 2 and 3) were collected during necropsy. Tissue was fixed for at least 48 hours in 4% buffered formaldehyde, embedded in paraffin, cut into 4 µm thick slides and stained with haematoxylin and eosin (H&E) following routine protocols [20]. Light microscopy was performed using the digital slide scanning system and ndp.view2 software, version 2.9.29 (both Hamamatsu photonics, Herrsching, Germany). H&E-stained sections were evaluated and lesions described. Scoring of lesions followed a semiquantitative scheme of four degrees: 1: minimal (<5 % of the area are affected or 1-3 foci), 2: mild (6-40% or >3 foci), 3: moderate (41-80%, or coalescing), 4: severe (>80%, or diffuse). A total lesion score for the CNS was calculated by cumulating individual lesion scores. Evaluation and interpretation were performed by a trained pathologist (LMM) and reviewed by a board-certified pathologist (DiplECVP, AB) in a masked fashion using the post-examination masking method [21].

### RNA *in situ* hybridization

Consecutive tissue slides from selected i.n./p.o. infected wood mice and rats from Exp. 3 were analysed by RNA-ISH as proof of principle analysis for target cell identification. RNA-ISH was performed using the RNAScope 2-5 HD Reagent Kit Red (Advanced Cell Diagnostics, Hayward, USA) according to the manufacturer’s instructions. A previously published custom-designed probe targeting the consensus sequence of a conserved stretch of the RusV nsPP open reading frame of the Swedish RusV clade 2 (genome pos. 66 to 427; catalogue no. 1145591-C1) was used to visualize RusV RNA [5]. Probes against the genes of peptidylprolyl isomerase B (PPIB) and dihydrodipicolinate reductase (DapB) served as technical controls. Tissues of non-infected wood mice and rats were used as negative controls. Target cells were identified based on their morphology.

### Statistical analysis

Statistical analysis of body weights determined in Exp. 2 or 3 was performed by Student’s t-test or One Way Analysis of Variance (ANOVA) with subsequent Tukeýs comparison of means, respectively, using GraphPad Prism version 8.4.3 (GraphPad Software, Boston, USA).

## RESULTS

### Intracranial inoculation of wood mice with RusV-positive brain homogenates

In Exp. 1, seven groups of four wood mice each were intracranially inoculated with RusV-positive brain homogenates originating from naturally infected *Apodemus* mice, cats or zoo animals originating from Sweden (groups A and D), Germany (groups B, C and E), Slovakia (group F) and Austria (group G). In all groups at least one animal had RusV RNA detectable by RT-qPCR in its brain, with all four animals being positive in three groups (groups B, D and E; Figure 1A). The correct identity of the virus was confirmed by Sanger sequencing of a 409 nt sequence stretch as described previously [5]. For all positive animals, the resulting sequence was identical to the respective inoculum (data not shown).

**Figure 1.**
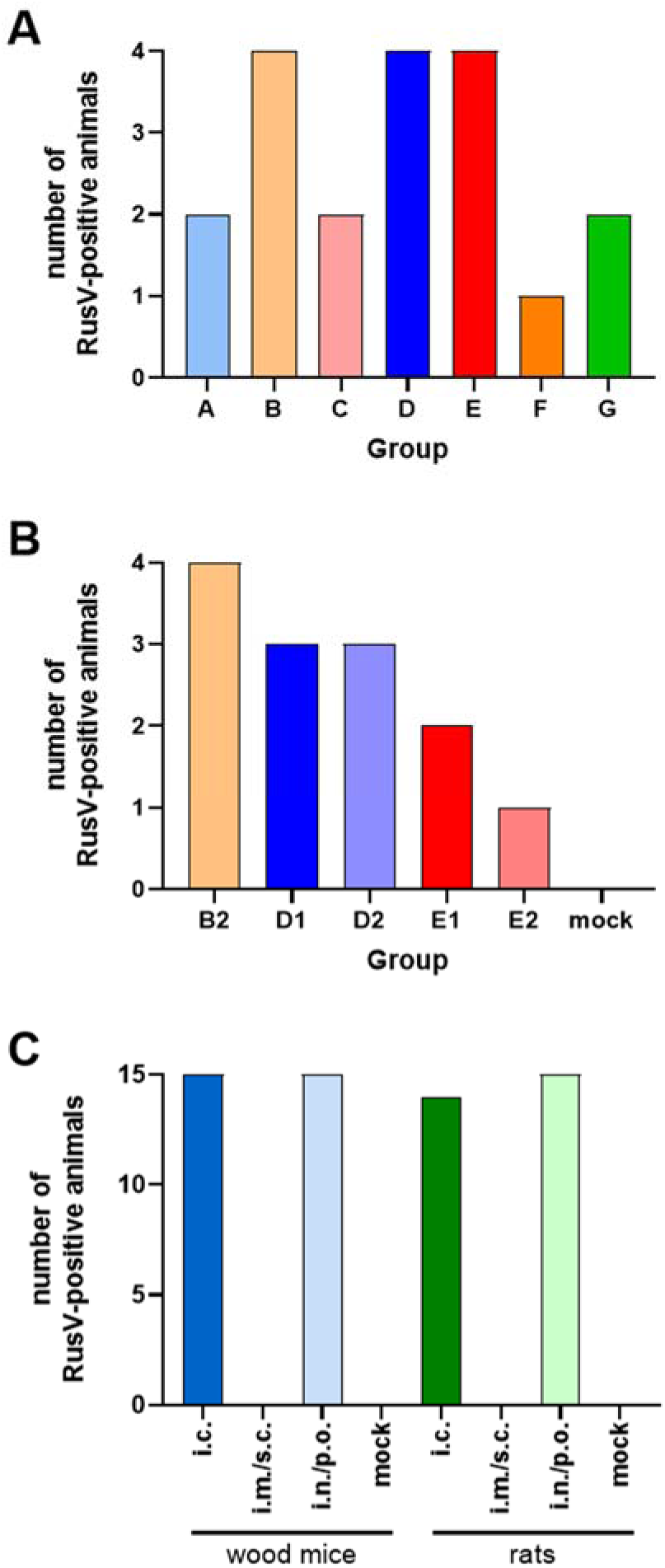
Frequency of RusV detection in rats and wood mice experimentally inoculated with brain homogenates of RusV-infected animals (Exp. 1 to 3). (A) Exp. 1: wood mice were inoculated intracranially (i.c.) with RusV-positive brain homogenates of seven different naturally infected animals. (B) Exp. 2: Lewis rats were i.c. inoculated with RusV-positive brain homogenates from either naturally infected animals or experimentally inoculated wood mice from Exp. 1. (C) Exp. 3: wood mice or Lewis rats were inoculated via the indicated routes with a pool of RusV-positive brain homogenates from experimentally inoculated rats from groups D1 and D2 of Exp. 2. Mock-inoculated animals received a brain homogenate of RusV-negative rats. An animal was considered positive if at least one of its tissues tested positive by RusV-specific RT-qPCR. i.m. = intramuscular; i.n. = intranasal; p.o. = peroral; s.c. = subcutaneous.

RT-qPCR Cq values in brain samples ranged from 22 to 31, with the lowest values observed in groups A and D inoculated with material of clade 2 from Sweden (Figure 2A). The animals from these two groups were also the only animals with low RusV RNA loads (Cq values 32 to 37) detectable in peripheral organs, such as intestine, lung, heart and kidney, with the sole exception of a single animal from group B also having very low viral loads detectable in its kidney (Figure 2A).

**Figure 2.**
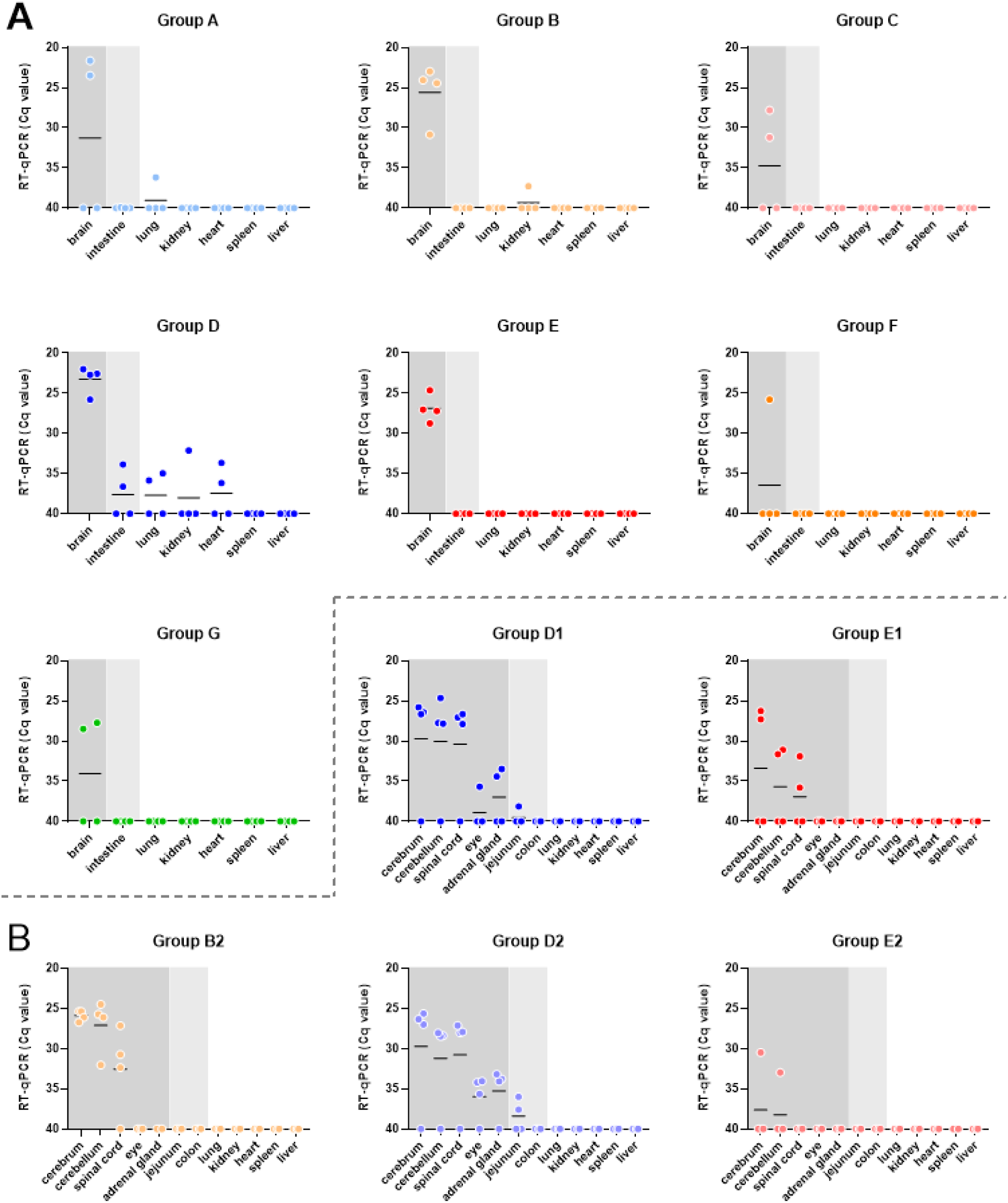
Tissue distribution of viral RNA in RusV-inoculated wood mice and rats (Exp. 1 & 2). (A) Exp. 1: wood mice were inoculated intracranially (i.c.) with RusV-positive brain homogenates of seven different naturally infected animals. (B) Exp. 2: Lewis rats were i.c. inoculated with RusV-positive brain homogenates from either naturally infected animals or experimentally inoculated wood mice from Exp. 1. Animals were euthanized at four weeks post inoculation. Tissue samples were tested for RusV-specific RNA by RT-qPCR. Results are presented as cycle of quantification (Cq) values. The position of the X axis indicates the detection limit of the assay. Organs are grouped into predominantly neural (dark grey) or mucosal tissues (light grey) and large parenchymal organs (white).

Weekly collection of swab samples from all animals resulted in only one sample with low levels of detectable RusV RNA (Cq 36). This sample had been collected from a RusV-positive animal of group A in week 3 p.i. (data not shown). No viral RNA was detected in the pooled faecal samples collected from each group at weekly intervals (data not shown).

RusV-reactive antibodies were not detectable by IFAT in any of the sera collected at euthanasia of the animals (data not shown).

Eight animals from three different groups (D, E and F) showed ruffled fur and weight loss during the course of the study and seven of them had to be euthanized at 15 to 26 days p.i. after reaching the termination criteria of >20% weight loss (Supplemental Fig. S1). The affected animals comprised RusV-positive as well as negative mice. None of the animals exhibited neurological signs. Histopathology was not performed in this experiment.

### Intracranial inoculation of Lewis rats with RusV-positive brain homogenates

In Exp. 2, five groups of four Lewis rats were inoculated with RusV-positive brain homogenates originating from either naturally infected animals or experimentally infected wood mice from the previous experiment (Exp. 1). Rats of groups D1 and E1 received the same inoculum from naturally infected animals as the wood mice of groups D or E, respectively of Exp. 1. Groups B2, D2 and E2 received pooled brain homogenates from the RusV-positive animals of groups B, D or E, respectively of Exp. 1 (Supplemental Table S2). An additional group of two animals remained uninfected and served as controls.

In all inoculated groups at least one animal had RusV RNA detectable in the brain (Figure 1B). Highest viral levels were detected in the cerebrum and cerebellum, followed by spinal cord. The Cq values in the CNS were overall higher than in the wood mice of Exp. 1 (Cq 24 to 30). Low viral RNA levels were also detectable in eye, adrenal gland and intestine of some RusV-positive animals of groups D1 and D2, which had received viral material originating from a Swedish cat (Figure 2B). Viral RNA was not detected in any oral swab or pooled faecal sample collected at weekly intervals during the experiment (data not shown).

None of the rat sera collected at the end of the experiment exhibited detectable reactivity in the RusV IFAT (data not shown).

None of the animals showed detectable clinical signs. However, at the end of the experiment at week 4 p.i., RusV-positive rats showed a tendency of reduced weight gain as compared to RusV-negative rats (*P* = 0.128; Student’s t-test; Supplemental Figure S2B).

Histological analysis revealed findings compatible with non-suppurative meningoencephalomyelitis in ten out of the 13 RusV-positive rats, either as individual findings or as a combination of several changes. These included perivascular accumulation of lymphocytes and macrophages (“perivascular infiltrates”), partially with activation of vascular endothelium or cell death within the perivascular space (“perivascular inflammation”), increased numbers of rod-shaped microglial cells (“microgliosis”), neuronal and glial single cell apoptosis/necrosis found disseminated in the brain and in some animals also prominently affecting the hippocampus (Figure 3A-D). Animals infected with the Swedish strain NRL.22_007-6 (groups D1 and D2) achieved higher cumulative lesion scores (scores 5 to 17) than those infected with other brain homogenates (scores 0 to 3; Figure 4A, Supplemental Figure S3). Only one out of nine RusV-negative rats exhibited minimal microgliosis in the brain and single-cell necrosis in the spinal cord. In peripheral organs, minimal to mild changes were documented in several animals, without apparent difference between RusV-positive and -negative rats (Supplemental Table S4).

**Figure 3.**
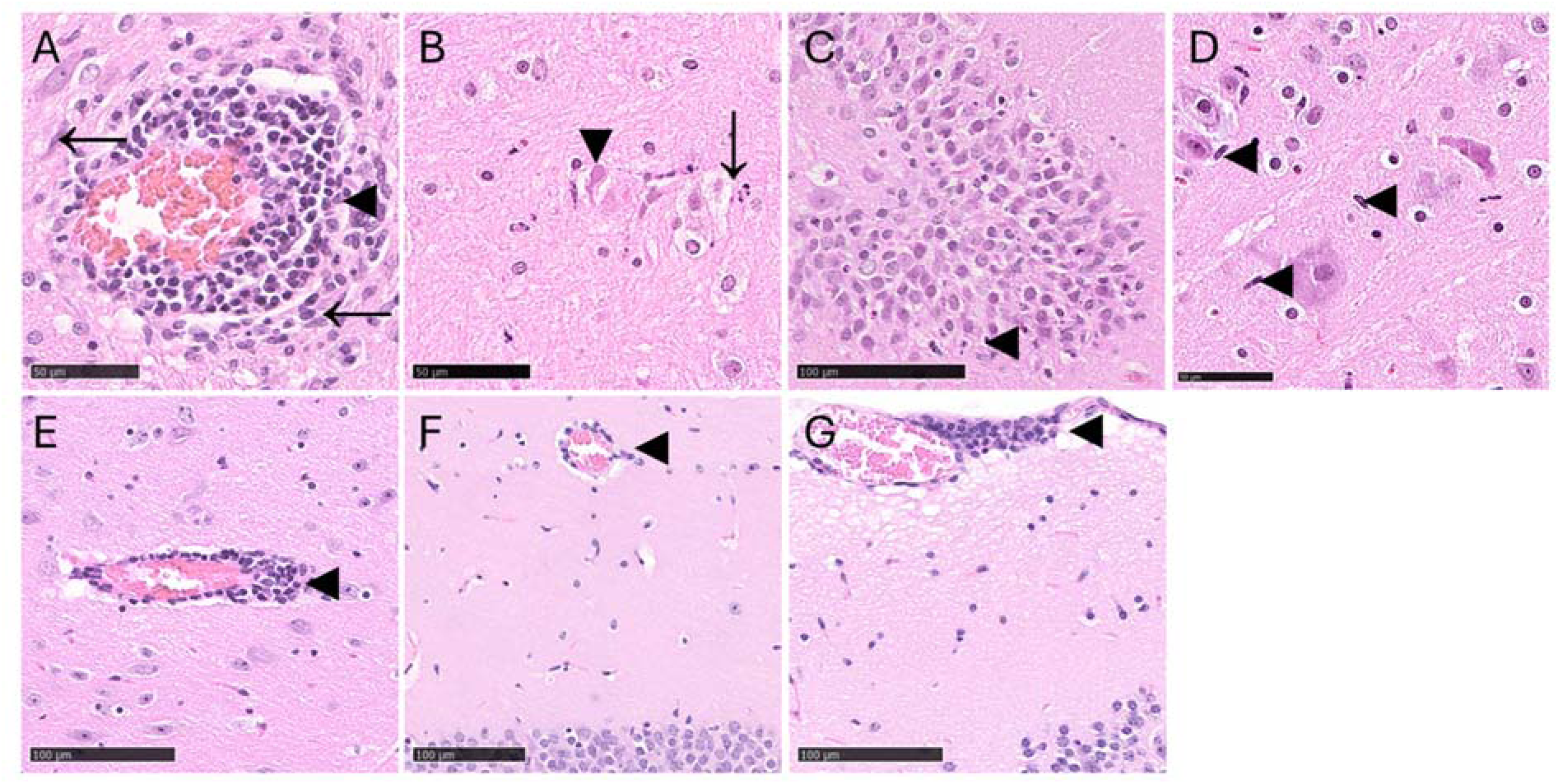
Histopathology in the brains of i.c.-inoculated RusV-positive rats (A-D; Exp. 2 & 3) and wood mice (E-G; Exp. 3). (E-G) Wood mice: Representative examples of perivascular lymphohistiocytic infiltrates (“perivascular cuffing”◄) in the thalamus (E), hippocampus (F), and in the meninges (G). (A-D) Rats: Perivascular, lymphohistiocytic infiltrates (◄) with associated tissue reaction (inflammation, ←) (A), necrosis of glial cells (←) and neurons (◄) (B), also affecting neurons (◄) in the hippocampus (C), increased number of rod-shaped microglial cells (microgliosis, ◄) (D). Hematoxylin-Eosin stain. Bars represent 50 or 100 µm, as indicated.

**Figure 4.**
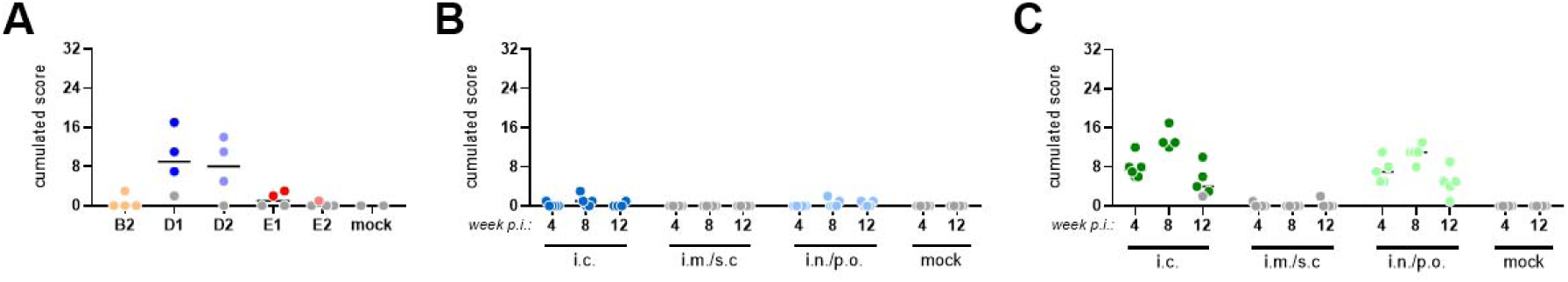
Cumulative microscopic lesion score in the central nervous system of RusV-inoculated rats and wood mice (Exp. 2 & 3). **(A)** Juvenile Lewis rats were i.c. inoculated with RusV-positive brain homogenates from either naturally infected animals or experimentally inoculated wood mice from Exp. 1. All animals were euthanized at four weeks post infection. **(B, C)** Adult wood mice **(B)** or juvenile Lewis rats **(C)** were RusV-inoculated via the indicated routes. The presence and severity of microscopic lesions was semiquantitatively scored by blinded evaluation. Results are presented as cumulated scores of eight individual scores representing perivascular infiltrates or inflammation, microgliosis and single cell necrosis/apoptosis in brain and spinal cord (Supplemental Figures S3, S5 and S6). Coloured dots represent RusV-positive animals, while grey dots represent RusV-negative animals, as determined by RT-qPCR. i.c. = intracranial; i.m./s.c. = intramuscular/subcutaneous; i.n./p.o. = intranasal/peroral.

### RusV inoculation of wood mice and Lewis rats via different routes

In Exp. 3, three groups of 15 mice and Lewis rats each were inoculated with brain homogenate originating from experimentally RusV-infected wood mice and rats originating from groups D, D1 and D2 of Exp. 1 and 2. The original inoculum of these groups derived from cat NRL.22_007-06 from Sweden. Groups A (wood mice) and E (rats) were injected intracranially. Groups B (wood mice) and F (rats) received a combined intramuscular and subcutaneous inoculation, while the inoculum was applied intranasally and perorally to groups C (wood mice) and G (rats). Two further groups (D, wood mice and H, rats) remained uninfected. After 4, 8 and 12 weeks p.i., five animals per group were euthanized.

With the exception of one i.c.-inoculated rat, euthanized at 12 weeks p.i., all i.c. and i.n./p.o.-inoculated animals of both species had RusV RNA detectable in their CNS. In contrast, RusV RNA was not detectable in the organs of any of the i.m./s.c.-inoculated wood mice or rats. Likewise, all mock-inoculated animals remained RusV-negative (Figure 1C).

Wood mice infected via i.c. or i.n./p.o. inoculation showed comparable RusV distribution profiles at all three time points. Viral RNA loads were highest in the CNS and other neuronal tissues (minimal Cq values 20 to 22), but viral RNA was consistently detectable also in mucosal organs and associated exocrine glands such as nose, salivary gland, intestine and urinary bladder (Cq ≥25). Only low RNA levels (Cq ≥31) were sporadically detected in heart, lung, spleen and kidney, but not in the liver (Figure 5A, B). The viral distribution was likewise comparable among i.c.- or i.n./p.o.-inoculated rats, but viral RNA levels were overall lower as compared to the wood mice. Cq values in the CNS reached only approximately 25 in both groups. Viral RNA was inconsistently detected in mucosal organs and the levels remained lower than in the wood mice (Cq ≥30). Low levels of RusV RNA (Cq ≥31) were also detected in the heart, lung, spleen and kidney of few i.n./p.o.-inoculated rats, but not in the i.c.-inoculated group (Figure 5C, D).

**Figure 5.**
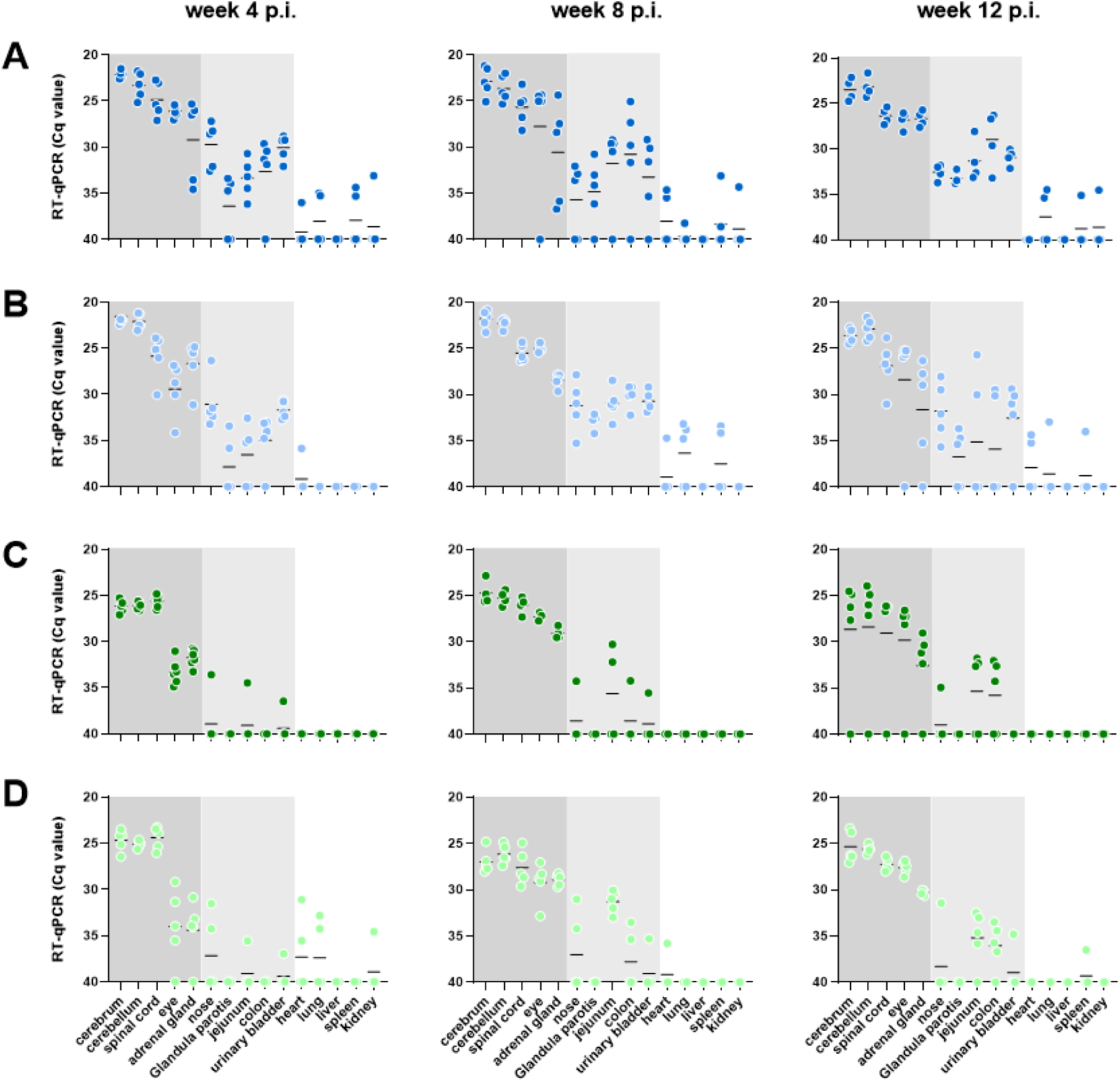
Tissue distribution of viral RNA in RusV-inoculated wood mice and rats (Exp. 3). Wood mice and Lewis rats were inoculated via different routes with a pool of RusV-positive brain homogenates from experimentally inoculated rats from groups D1 and D2 of Exp. 2. Five animals per group were euthanized at four, eight or twelve weeks post inoculation (p.i.). Tissue samples were tested for RusV-specific RNA by RT-qPCR. Results are presented as cycle of quantification (Cq) values. The position of the X axis indicates the detection limit of the assay. Organs are grouped into predominantly neural (dark grey) or mucosal tissues (light grey) and large parenchymal organs (white). Only intracranially (i.c.) infected wood mice (A), intranasally (i.n.)/perorally (p.o.) inoculated wood mice (B), i.c. inoculated rats (C) and i.n./p.o. inoculated rats (D) are shown. All examined organs of intramuscularly/subcutaneously or mock-inoculated wood mice and rats tested negative (data not shown).

Oral swabs were collected at weekly intervals and analysed using an optimized protocol as compared to Exp. 1 and 2. With the exception of one RusV-positive swab collected already at week 1 p.i., infected wood mice started shedding detectable amounts of viral RNA at 3 weeks p.i. and remained at constant but relatively low levels (Cq ≥29) from week 4 p.i. to the end of the experiment.

Detection rates in the two RusV-positive groups ranged from 40 to 100 % during this period (Figure 6A). Very low RusV RNA loads (Cq ≥34) were also detected in up to two i.n./p.o.-inoculated rats from 5 to 11 weeks p.i., whereas all swabs collected from i.c.-inoculated rats tested negative (Figure 6A). Environmental swabs were collected from the cage walls from each subgroup of 5 animals. Between weeks 5 to 11 p.i. very low levels of viral RNA (Cq ≥34) could be detected in environmental samples collected from infected wood mice with no apparent difference between the two positive groups, but not from RusV-positive rats (Figure 6B). Pooled faecal samples tested very weakly positive (Cq ≥35) only for two different cages of i.n./p.o.-inoculated wood mice at weeks 2 and 6 p.i. (Figure 6C).

**Figure 6.**
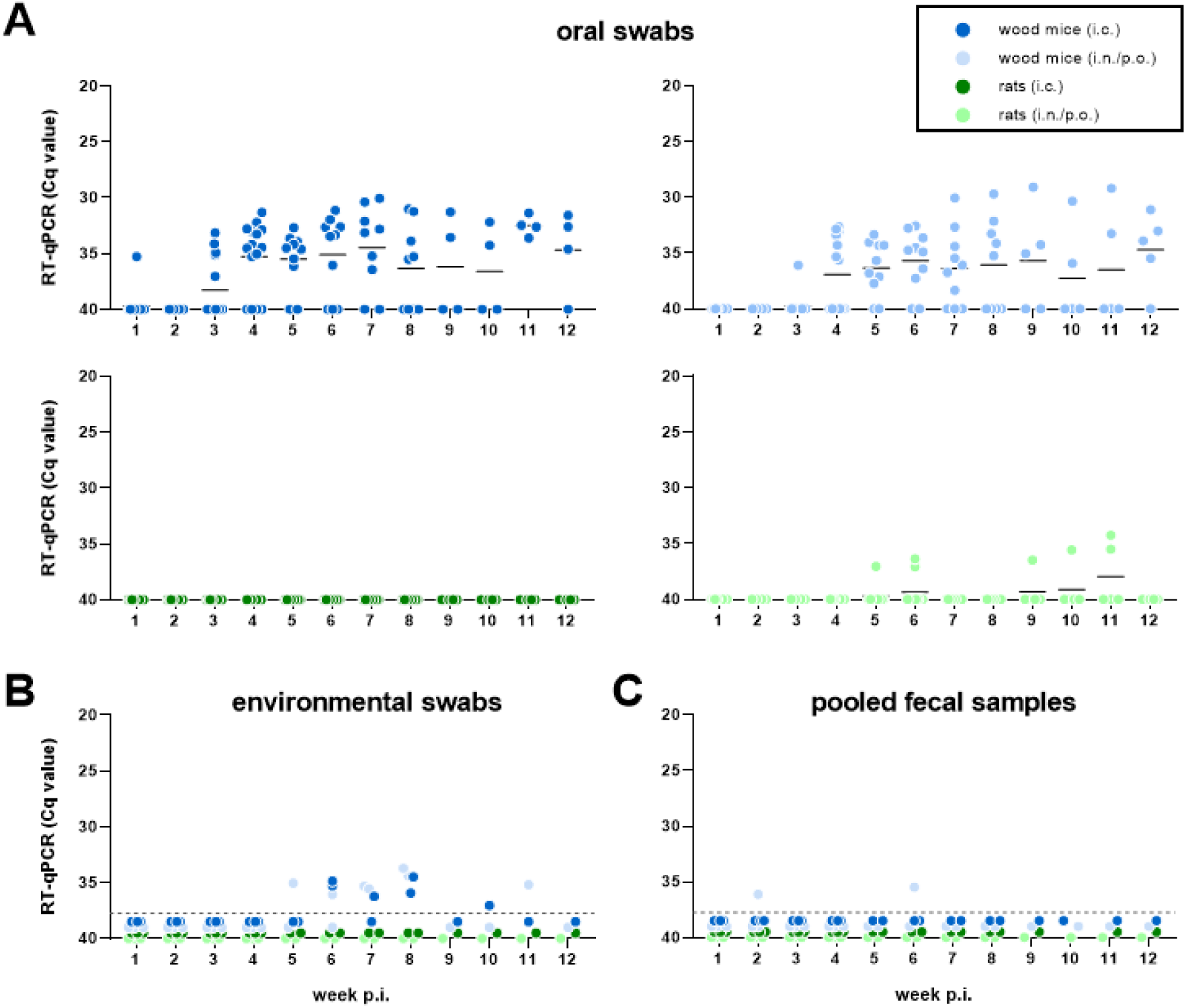
Detection of viral RNA in sheddings of RusV-inoculated wood mice and rats (Exp. 3). Wood mice and Lewis rats were inoculated via the indicated routes with a pool of RusV-positive brain homogenates from experimentally inoculated rats from groups D1 and D2 of Exp. 2. (A) Oral swabs were collected from each individual at weekly intervals. (B, C) Environmental swabs taken from the cage walls (B) and pooled faecal samples (C) were collected at weekly intervals from each cage of five animals. All samples were tested for RusV-specific RNA by RT-qPCR. Results are presented as cycle of quantification (Cq) values. The position of the X axis indicates the detection limit of the assay. Dots below the dashed line indicate negative RT-qPCR results (no Cq). i.c. = intracranial; i.n./p.o. = intranasal/peroral; p.i. = post inoculation.

The presence of serum antibodies, as detected by IFAT, was observed only in five i.n./p.o.-inoculated wood mice euthanized at 8 or 12 weeks p.i., with IFAT titres ranging from 20 to 1,280 (Supplemental Figure S4A). Three of the sera showed reactivity only with the E2/E1 protein and one only with the capsid protein. Only one serum reacted with both antigens (Supplemental Figure S4B).

None of the animals of both species exhibited neurological signs during the course of the experiment. However, one i.c.-inoculated rat died during anaesthesia in week 4 p.i. and one i.c.-inoculated wood mice had to be euthanized prematurely at week 6 p.i. due to progressing weight loss. Body weights of the wood mice showed a considerable variance, making assessments of potential RusV-related differences difficult (Figure 7). Rats of the i.c.- and i.n./p.o.-inoculated groups started to significantly gain less weight as compared to the RusV-negative i.m./s.c.- and mock-inoculated groups in weeks 5 to 6 p.i. (One Way ANOVA with subsequent Tukeýs comparison of means; *P*<0.05). At the end of the experiment they reached only about 83 % of the weight of the non-infected groups (Figure 7).

**Figure 7.**
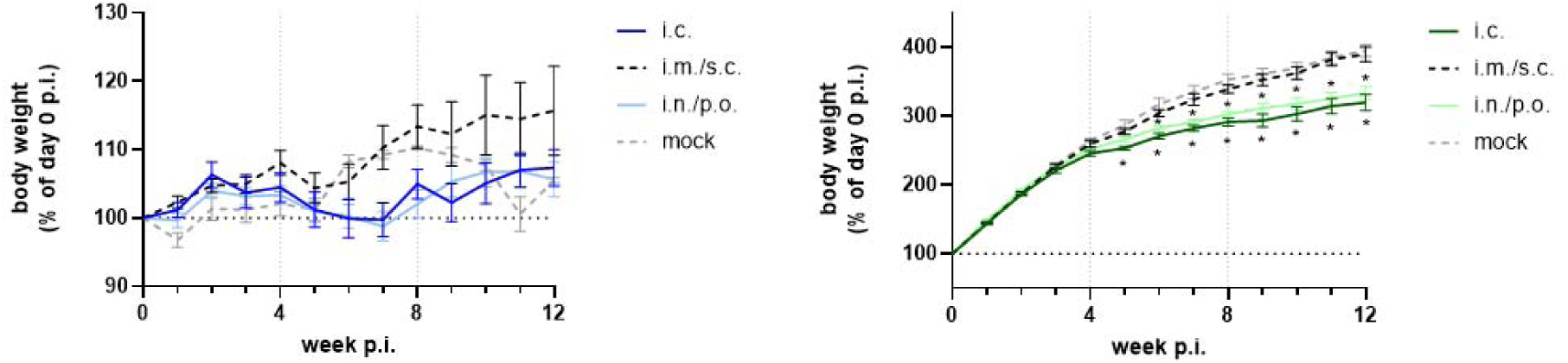
Body weight of RusV-inoculated wood mice and rats (Exp. 3). Adult wood mice and juvenile Lewis rats were inoculated via the indicated routes with a pool of RusV-positive brain homogenates from experimentally inoculated rats from groups D1 and D2 of Exp. 2. Body weights were determined at weekly intervals. Results are presented as arithmetic mean (+/- standard deviation) per group of the relative body weight (% of the body weight at the start of the experiment; day 0 post infection, p.i.). Solid coloured lines represent groups with RusV-positive animals, whereas broken grey or black lines represent groups with only RusV-negative animals as determined by RT-qPCR. i.c. = intracranial; i.m./s.c. = intramuscular/subcutaneous; i.n./p.o. = intranasal/peroral. Asterisks represent statistically significant differences to both uninfected groups (One Way ANOVA with subsequent Tukeýs comparison of means; *P*<0.05).

### Histopathology in rats and wood mice inoculated via different routes

Ten out of 30 RusV-positive wood mice, showed minimal to mild lesions in the brain, but not in the spinal cord, with cumulative lesion scores not exceeding 3. None of the RusV-negative wood mice were affected (Figure 3E-G, 4B, Supplemental Figure S6). The most frequent findings were perivascular infiltrates (n=7) and single cell necrosis (n=4), while microgliosis was not observed. With the exception of one i.c.-inoculated animal euthanized at week 4 p.i. and the i.c.-inoculated animal that had to be euthanized at week 6 p.i., only wood mice euthanized at 8 or 12 weeks p.i. exhibited lesions (Figure 4B, Supplemental Figure S6, Supplemental Table S5).

In contrast to the wood mice, RusV-infected rats exhibited more prominent meningoencephalomyelitis, covering the full spectrum of lesions (Figure 3A-C), with cumulative lesion scores of up to 17. Only three of the 26 RusV-negative rats reached lesion scores of up to 2 (Figure 4C, Supplemental Figure S6). No apparent differences were observed between the i.c. and the i.n./p.o.-inoculated rats and both groups showed a similar time course of lesion development. Perivascular infiltration and inflammation appeared to peak in week 8 p.i. as compared to weeks 4 and 12 p.i., whereas single cell necrosis scores were highest already at week 4 p.i. followed by a gradual decline thereafter (Figure 4C, Supplemental Figure S6).

Lesions in peripheral organs affecting RusV-positive as well as negative wood mice and rats are summarized in Supplemental Tables S5 and S6.

### RusV cell tropism in RusV-infected wood mice and rats

The viral distribution and target cells of RusV in the brain and spinal were analysed by RNA-ISH for selected i.n./p.o.-inoculated wood mice euthanized at 4 (n=3), 8 (n=2) or 12 (n=3) weeks p.i. as well as rats euthanized at 4 (n=4) and 12 (n=3) weeks p.i. of Exp. 3. In all analysed wood mice, RusV RNA was abundantly found in neurons throughout all analysed brain areas, in the grey matter of the spinal cord as well as in adjacent route nerves and dorsal root ganglia neurons (Supplemental Figure S7A). The fine-granular to globular signal was mainly located in the perikaryon of neurons, but also fine-granular in the neuropil and in the white matter (Supplemental Figure S8A-H). Cell borders in the brain cannot be reliably identified, thus it is impossible to exclude other target cells at present. As compared to wood mice, viral RNA was less abundant in the brains of seven analysed RusV-infected rats that had been euthanized at weeks 4 (n=4) or 12 (n=3) p.i. (Supplemental Figure S7B, S8I-L).

The cellular tropism was furthermore analysed by RNA-ISH in peripheral tissues for nine i.n./p.o.-inoculated wood mice euthanized in weeks 4 (n=4), 8 (n=1) and 12 (n=4) p.i. However, not all tissues were available for all of these animals (Supplemental Figure S7C). RusV RNA was detected not only in neurons of the optical nerve, the retina (ganglion cell layer, inner plexiform layer, inner nuclear layer) and intramural ganglia of the stomach and intestine, but also in neuroendocrine epithelial cells of the pancreas islets and adrenal medulla as well as in the glandular epithelium of maxillary sinus glands of individual wood mice (Supplemental Figure S7C, S9). No RusV-positive cells were detectable in samples of the heart, liver, spleen, salivary glands with lymph nodes, lung, kidney or urinary bladder (data not shown).

## DISCUSSION

In this study we successfully established first *in vivo* infection models for RusV and collected important insights into infection routes, course of infection, tissue and cell tropism and possible routes of viral shedding. Furthermore, we could reproduce RusV-associated meningoencephalomyelitis in the rat model.

Using brain homogenates from naturally and later also experimentally infected animals, inoculation of wood mice and rats was successful via i.c. injection and also by a combined mucosal route (i.n./p.o.), which is presumably representing the natural route of infection. Whether RusV may enter the organism via i.n., p.o. or both routes equally will need to be determined in follow-up experiments. Surprisingly, RusV failed to establish infection when administered parenterally (i.m./s.c.) in both examined species.

Brain homogenates had to be used for inoculation since isolation of the virus in cell culture had failed so far despite employing various primary and established cell lines and different cultivation procedures. Attempts to isolate the virus using fresh brain tissue originating from the experiments of this study likewise failed, even when preparing primary brain cell cultures from infected animals, followed by cocultivation with various infinite cell lines (unpublished observations). In contrast, i.c. inoculation of wood mice led to infection for all selected brain samples from naturally infected animals, including even a brain tissue pool from Austrian cats from the early 1990s, which had been thawed and refrozen several times meanwhile. While a standardized comparison of the performance of the different viruses was impossible due to the lack of a virus titration method, the viruses representing RusV clade 2 from Sweden appeared to have the tendency of a more rapid spread from the CNS into the periphery in both, wood mice and rats, and it induced more prominent encephalitis in infected rats. Thus, we selected ‘isolate’ NRL.22_007-6 (genotype 2A from a Swedish cat) as the inoculum for the most extensive experiment 3.

As described previously for naturally infected yellow-necked field mice and encephalitic cats and zoo animals [1, 3, 4, 9, 16], highest loads of viral RNA were consistently found in the CNS, with only minor differences between cerebrum and cerebellum and slightly lower levels detectable in the spinal cord in both species tested. Viral RNA was also consistently detectable in further organs containing considerable proportions of neuronal tissue, such as eye and adrenal gland. Mean Cq values in the brains of wood mice and rats differed by about 3, representing approximately 10-fold higher RNA loads in the wood mice. This was also confirmed by RNA-ISH, which demonstrated RusV RNA to be widespread and homogenously distributed in the brains of wood mice, while the staining was less abundant in rats. RNA-ISH identified neurons as the predominant target cells in the brain, spinal cord, optic nerve, and retina and occasionally also in the peripheral nervous system, including dorsal route ganglia and root nerves of the spinal cord, confirming pervious observations from naturally infected animals [1, 3–7, 9, 16].

Whether also glial cells are infected requires claification by double labelling approaches in future studies. In addition, viral RNA was also detected in neuroendocrine epithelial cells of the adrenal gland medulla of infected wood mice, as already described for RusV-infected yellow-necked field mice [16], as well as of the pancreas.

RT-qPCR detected low to moderate levels in various peripheral tissues of wood mice, including several containing or having access to mucosal surfaces, such as nose, salivary glands, intestine and urinary bladder. Only very low levels of viral RNA were sporadically detected in heart, lung, kidney and spleen of wood mice, while the liver remained negative throughout the study. RNA-ISH of selected wood mouse tissues confirmed RusV RNA not only in intramural ganglion neurons of the stomach and intestine as part of the peripheral nervous system, but also in epithelial cells of the paranasal maxillary sinus gland. The tissue distribution in rats followed the same general pattern as in wood mice, but viral loads and detection frequencies were markedly lower, which is consistent with the lower viral RNA levels in their CNS. In wood mice, viral loads and tissue distribution patterns did not differ between the necropsies at 4, 8 and 12 weeks p.i., indicating a long-term, possibly life-long, persistence of the virus. In rats, the only prominent difference was a marked increase in viral loads in eye and adrenal gland from week 4 to 8 p.i., demonstrating a somewhat slower viral kinetic as compared to the wood mice. In both species, the distribution patterns did not differ between the i.c. or i.n./p.o. inoculation route. It may be speculated that the virus may directly enter the brain following mucosal inoculation, e.g. via the olfactory nerve or other cranial nerves.

The consistent detection of RusV RNA in mucosal tissues of wood mice infected with RusV of clade 2 is in contrast to yellow-necked field mice from northeastern Germany, which were naturally infected with RusV of clade 1. In the latter, viral RNA had been detectable almost exclusively in the CNS [1, 16]. Whether this discrepancy is associated with the host species, the viral strain, the time point after infection or combinations of them remains to be elucidated. In congruence with the presence of the virus in nose and salivary glands, low to moderate levels of viral RNA could be detected in oral swabs collected from the experimentally infected wood mice starting around week 4 p.i. Very low viral RNA levels were also detectable in faeces and in environmental swabs from cage walls. The viral RNA might originate from mucus or saliva, but also from unsampled sources, such as urine. Due to the lack of a virus isolation method for RusV, it remains unknown if the detected viral RNA in sheddings represents infectious virus. Likewise, the way by which RusV RNA is shed from mucosal surfaces of infected wood mice remains to be determined, but detection of viral RNA in the maxillary sinus gland epithelium suggests a potential secretion via mucus. A structured examination of the mucus-producing glands of the gastrointestinal and upper respiratory tract is mandatory for future studies. In agreement with the sparse presence of viral RNA in their peripheral organs, oral swabs of rats were barely positive and only for the i.n./p.o.-inoculated group. Our results suggest rats to rather not act as a RusV reservoir host.

Surprisingly, seroconversion was detected by IFAT only in few RusV-infected wood mice inoculated by the mucosal route, but not in rats or i.c.-inoculated wood mice. Reactivities were observed against both, the capsid and the E2/E1 protein with usually rather low titres. It remains to be elucidated whether this infrequent detection of RusV-reactive antibodies is a result of a rather low sensitivity of the established IFAT or whether seroconversion during RusV infection is indeed rare. Alternative serological tests for RusV or attempts to detect RusV-reactive antibodies in naturally infected animals have not been reported, yet.

Histopathological analysis revealed findings consistent with a mostly mild to moderate non-suppurative meningoencephalomyelitis in RusV-infected rats, including perivascular lymphohistiocytic infiltrates and inflammation, microgliosis, and apoptosis/necrosis of neurons and glial cells. The nature of these lesions matches the findings described for naturally RusV-infected zoo animals and cats [1, 3–7, 9, 10]. Necrosis scores were highest already in week 4 p.i., while inflammatory lesions and infiltrates appeared to peak in week 8 p.i., and all parameters had somewhat declined at the end of the experiment at week 12 p.i. In contrast to the rats, only about one third of the RusV-infected wood mice presented with minimal lesions of the brain, usually affecting only a single parameter. Since these lesions were not observed in RusV-negative wood mice in the same experiment, they appear to be associated with RusV infection. These findings contrast with the absence of detectable encephalitis in the brains of naturally RusV-infected yellow-necked field mice and wood mice, which might be explained by the often poor tissue quality of the naturally infected animals, which usually had been trapped or found dead [1, 3, 5, 16]. Furthermore, if RusV-induced inflammation was indeed reduced after a transient peak, as suggested for the experimentally infected rats, the unknown, but possibly longer duration of the infection may have contributed to the absence of detectable lesions in the naturally infected rodents. However, this must be interpreted with caution since the course of the infection at even later stages in wood mice was not addressed in the present study. Changes in peripheral organs were not associated with the infection status of the animals in both species and are thus considered as background findings.

Despite the detectable CNS lesions, neither experimentally infected wood mice nor rats developed typical neurological disorders, as described for RusV-infected zoo animals or staggering disease in cats, until week 12 p.i. [1, 3–10]. However, systematic neurological tests to identify more subtle clinical changes were not performed in this study. In juvenile rats, RusV infection resulted in a significantly reduced weight gain starting from week 4 to 5 p.i., indicating that the RusV-induced encephalomyelitis was not purely subclinical. Significant body weight reduction was not observed for infected wood mice.

In summary, both, wood mice and rats, readily establish an apparently persistent RusV infection following mucosal inoculation, likely reflecting the natural infection route. Experimentally infected wood mice possess higher viral loads in the CNS and a broader tissue distribution, as compared to rats, and they are shedding viral RNA. They do not develop detectable clinical signs or weight loss and only minimal lesions in the CNS. This profile is compatible with wood mice serving as a natural reservoir of RusV. Infected rats show overall less virus in CNS and particularly in the periphery, with barely detectable shedding of viral RNA, indicating that they are unlikely to act as a reservoir host. The significantly reduced body weight and the reproduction of a lymphohistiocytic meningoencephalomyelitis in experimentally infected rats, which resembles the lesions described for naturally infected cats and zoo animals, represents an important step towards fulfilling Henle-Koch’s postulates for this recently discovered virus. The relatively mild lesions and the absence of detectable neurologic signs in infected rats may be a result of rats being zoologically closely related to wood mice and yellow-necked field mice, all being rodents of the subfamily Murinae. Thus, RusV may still be partly adapted to rats, sufficient to at least prevent a full-blown disease.

This study provides the basis and models for future research on RusV, targeting e.g. the routes of transmission by reservoir hosts, the routes of viral spread into and from the CNS, the pathogenesis of RusV-induced disease, the optimization of diagnostic tools, further attempts to isolate the virus and attempts to fully reproduce the disease in non-rodent hosts.

## ACKNOWLEDGEMENTS

The authors like to thank Kathrin Steffen, Weda Hoffman, Doreen Schulz, Robin Brandt, Silvia Schuparis, Ole Pietsch, Philip Starcky and Patrick Zitzow for their excellent technical assistance. For excellent care of animals and support during trials we thank the animal caretakers Kerstin Kerstel, Tanja Janke, Nicole Sinkwitz and Felix Zimak. We are grateful to Dirk Höper for performing RIEMS analyses.

## AUTHOR CONTRIBUTIONS

Conception and design: DR, KS, AK, AB, MB.

Establishing diagnostic methods: JG, AE, DR

Performance of experiments and analyses: AK, LMM, CW, SCW, LU, AB, KS, DR.

Analysis and interpretation of the data: DR, AB, LMM, CW, SCW

Drafting of the paper: DR, AB.

Revising the paper critically for intellectual content: all authors.

All authors approve of the final version of the manuscript.

