## Supplemental Material for "*In vivo* models for rustrela virus (RusV) infection in potential reservoir and spill-over hosts"

**Supplemental Table S1. Experimental design and inocula of Exp. 1 (wood mice, intracranially inoculated).**

| Group | No. of animals | Brain homogenate used for inoculation | | | | | | | |
|---|---|---|---|---|---|---|---|---|---|
| | | **Sample** | **Host** | **country** | **year** | **RT-qPCR Cq value [a]** | **RusV genotype [b]** | **GenBank accession** | **Reference** |
| **A** | 4 | KS21/1362 | wood mouse (*Apodemus sylvaticus*) | Sweden | 2011 | 22.5 | 2B | ON641049.1 | [1] |
| **B** | 4 | BRA_220 | yellow-necked field mouse (*Apodemus flavicollis*) | Slovakia | 2014 | 21.0 | 1E | - | - |
| **C** | 4 | KS20/1512 | yellow-necked field mouse (*Apodemus flavicollis*) | Germany | 2020 | 18.9 | 1A | OL960720.1 | [2] |
| **D** | 4 | NRL.22_007-6 (SWE-15) | domestic cat (*Felis catus*) | Sweden | 2021 | 20.2 | 2A | ON641046.1 | [1] |
| **E** | 4 | NRL.20_131-7 | South American coati (*Nasua nasua*) | Germany | 2020 | 18.9 | 1A | OL960717.1 | [2] |
| **F** | 4 | NRL.21_105 | red-necked wallaby (*Notamacropus rufogriseus*) | Germany | 2021 | 23.2 | 1B | OP221677.1 | [3] |
| **G** | 4 | NRL.22_003 (Pool) | domestic cat (*Felis catus*) | Austria | 1992-1993 | 28.5 | 3A | ON641041.1 ON641042.1 | [1] |

a RusV RNA detected by RusV-specific RT-qPCR; shown as cycle of quantification (Cq) value.

b RusV genotype according to the nomenclature established by Pfaff et al. [4].

**Supplemental Table S2. Experimental design and inocula of Exp. 2 (Lewis rats, intracranially inoculated).**

| Group | No. of animals | Brain homogenate used for inoculation | | | | | | |
|---|---|---|---|---|---|---|---|---|
| | | **Sample** | **Host** | **country** | **year** | **RT-qPCR Cq value** | **RusV genotype** | **GenBank accession** |
| **B2** | 4 | pool of brains of RusV-positive wood mice of group B, Exp. 1; inoculated with sample BRA_220 | | | | | | |
| **D1** | 4 | NRL.22_007-6 | domestic cat (*Felis catus*) | Sweden | 2021 | 20.2 | 2A | ON641046.1 |
| **D2** | 4 | pool of brains of RusV-positive wood mice of group D, Exp. 1; inoculated with sample NRL.22_007-6 | | | | | | |
| **E1** | 4 | NRL.20_131-7 | South American coati (*Nasua nasua*) | Germany | 2020 | 18.9 | 1A | OL960717.1 |
| **E2** | 4 | pool of brains of RusV-positive wood mice of group E, Exp. 1; inoculated with sample NRL.20_131-7 | | | | | | |
| **F** | 2 | not inoculated | | | | | | |

**Supplemental Table S3. Experimental design and inoculum of Exp. 3 (wood mice and rats, different routes).**

All animals were inoculated via different routes with pooled brain homogenates of all RusV-positive animals of groups D of Exp. 1 and D1 and D2 of Exp. 2, which had been inoculated with virus originating from the infected cat NRL.22_007-6, genotype 2A, from Sweden [1].

| Group | Species | Inoculation route [a] | No. of animals | No. of animals euthanized at week post inoculation | | | |
|---|---|---|---|---|---|---|---|
| | | | | 4 | 6 | 8 | 12 |
| A | wood mice | i.c. | 15 | 5 | 1 [b] | 5 | 4 [b] |
| B | wood mice | i.m./s.c. | 15 | 5 | - | 5 | 5 |
| C | wood mice | i.n./p.o. | 15 | 5 | - | 5 | 5 |
| D | wood mice | mock | 10 | 5 | - | - | 5 |
| E | rats | i.c. | 15 | 6 [c] | - | 4 [c] | 5 |
| F | rats | i.m./s.c. | 15 | 5 | - | 5 | 5 |
| G | rats | i.n./p.o. | 15 | 5 | - | 5 | 5 |
| H | rats | mock | 10 | 5 | - | - | 5 |

[a] i.c.: intracranial; i.m./s.c.: intramuscular/subcutaneous; i.n./p.o.: intranasal/peroral.

[b] One i.c.-inoculated wood mouse had to be euthanized prematurely in week 6 post inoculation (p.i.) due to progressive weight loss.

[c] One i.c.-inoculated rat died during anaesthesia for sample collection in week 4 p.i.

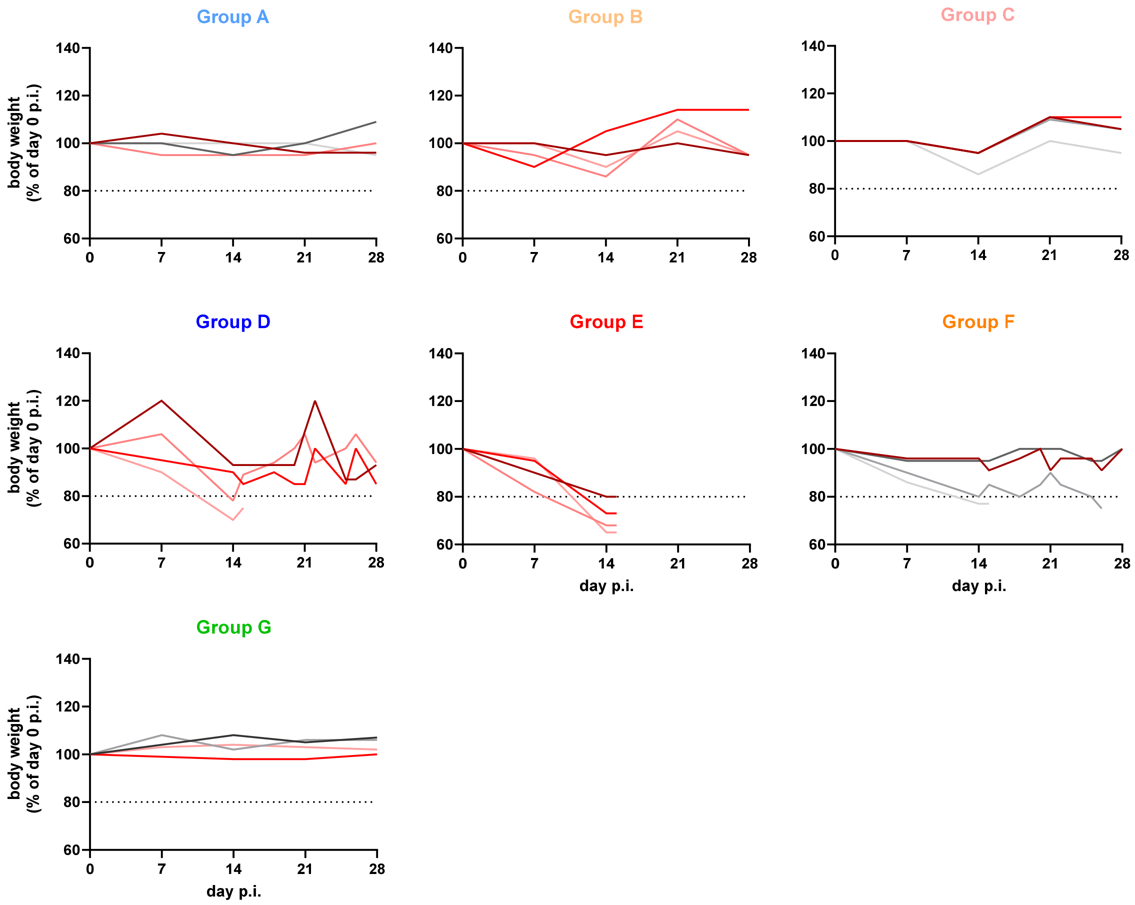


**Supplemental Figure S1. Body weight of RusV-inoculated wood mice (Exp. 1).** Wood mice were inoculated intracranially (i.c.) with RusV-positive brain homogenates of seven different naturally infected animals. Body weights were determined at weekly intervals or when animals showed clinical signs. Red lines indicate RusV-positive animals whereas grey lines indicate animals that tested RusV-negative. Results are presented as % of the body weight at the start of the experiment (day 0 post infection, p.i.). The dotted line represents the threshold at which animals were euthanized in line with the animal welfare criteria as approved by the competent authority.

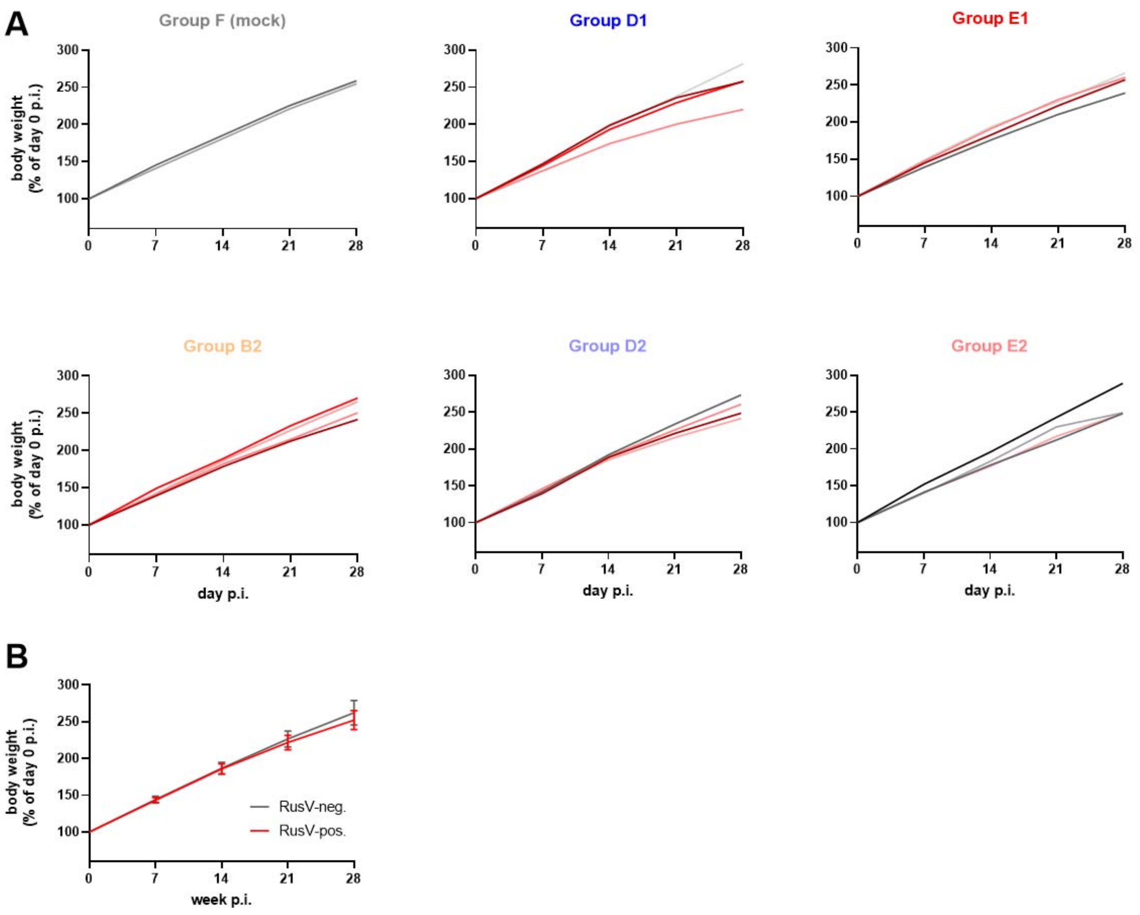


**Supplemental Figure S2. Body weight of RusV-inoculated rats (Exp. 2).** (A) Juvenile Lewis rats were intracranially (i.c.) inoculated with RusV-positive brain homogenates from either naturally infected animals or experimentally inoculated wood mice from Exp. 1. Body weights were determined at weekly intervals. Red lines indicate RusV-positive animals, whereas grey lines indicate animals that tested RusV-negative. Results are presented as % of the body weight at the start of the experiment (day 0 post infection, p.i.). (B) Arithmetic mean (+/- standard deviation) of relative body weights of all RusV-positive (red, n=13) and RusV-negative (grey, n=9) rats of Exp. 2.

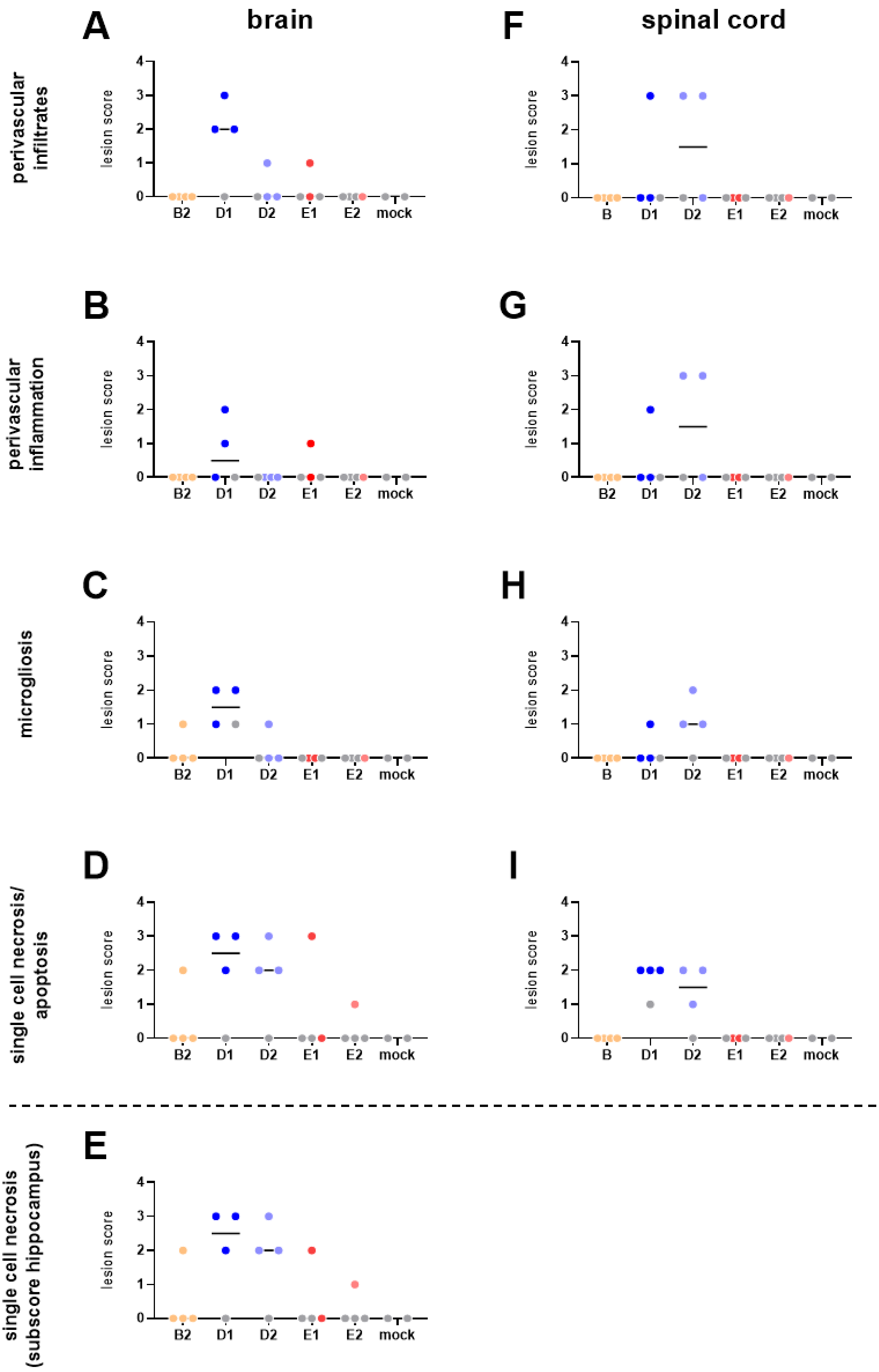
A
brain
F
spinal cord
perivascular infiltrates
B
G
perivascular inflammation
C
H
microgliosis
D
I
single cell necrosis/ apoptosis
E
single cell necrosis (subscore hippocampus)
lesion score
0
1
2
3
4
B2
B
D1
D2
E1
E2
mock

**Supplemental Figure S3. Individual microscopic lesion scores in the central nervous system of RusV-inoculated rats (Exp. 2).** Juvenile Lewis rats were intracranially (i.c.) inoculated with RusV-positive brain homogenates from either naturally infected animals or experimentally inoculated wood mice from Exp. 1. All animals were euthanized at four weeks post infection. The presence and severity of microscopic lesions was semiquantitatively scored by blinded evaluation. Score 0: lesion not detectable, 1: minimal (<5% of the area are affected or 1-3 foci), 2: mild (6%–40% or >3 foci), 3: moderate (41%–80%, or coalescing), 4: severe (>80%, or diffuse). Results are presented as individual scores for the indicated parameters in brain and spinal cord. Coloured dots represent RusV-positive animals, while grey dots represent RusV-negative animals, as determined by RT-qPCR. Raw data underlying this figure are provided as Supplementary Table S4.

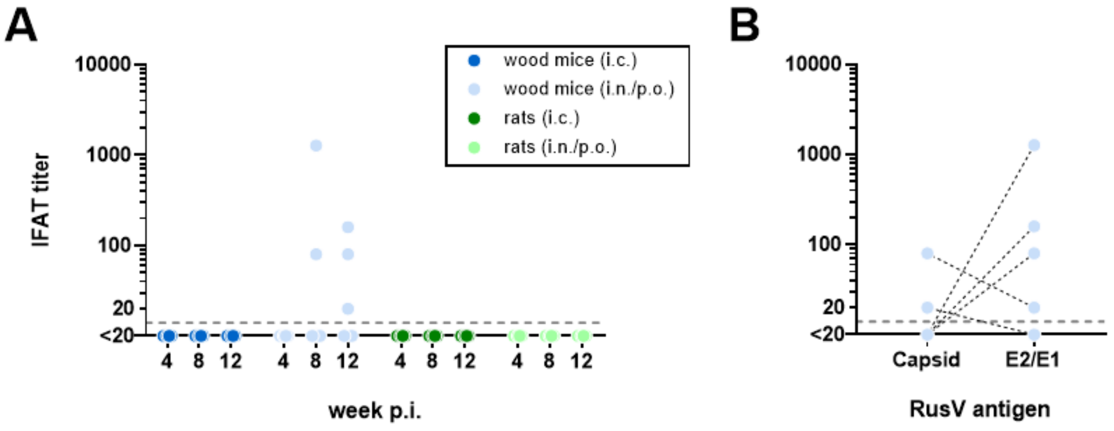


**Supplemental Figure S4. Seroconversion of RusV-inoculated wood mice and rats (Exp. 3).** Wood mice and Lewis rats were inoculated via the indicated routes with a pool of RusV-positive brain homogenates from experimentally inoculated rats from groups D1 and D2 of Exp. 2. Five animals per group were euthanized at four, eight or twelve weeks post inoculation (p.i.) and serum samples were collected. RusV-reactive antibodies were quantified using endpoint titration by immunofluorescence antibody test (IFAT) using transfected cells expressing RusV antigens. The position of the X axis indicates the detection limit of the assay. (A) Initial screening of all sera on cells co-expressing Capsid and E2/E1 proteins. Only results of the groups with detection of RusV RNA by RT-qPCR are shown. All examined sera of intramuscularly/subcutaneously or mock-inoculated wood mice and rats tested negative (data not shown). (B) Comparative titration of only the five positive sera on cells expressing either RusV capsid or E2/E1 protein. Dotted black lines connect results of the same sample. i.c. = intracranial; i.n./p.o. = intranasal/peroral; p.i. = post inoculation.

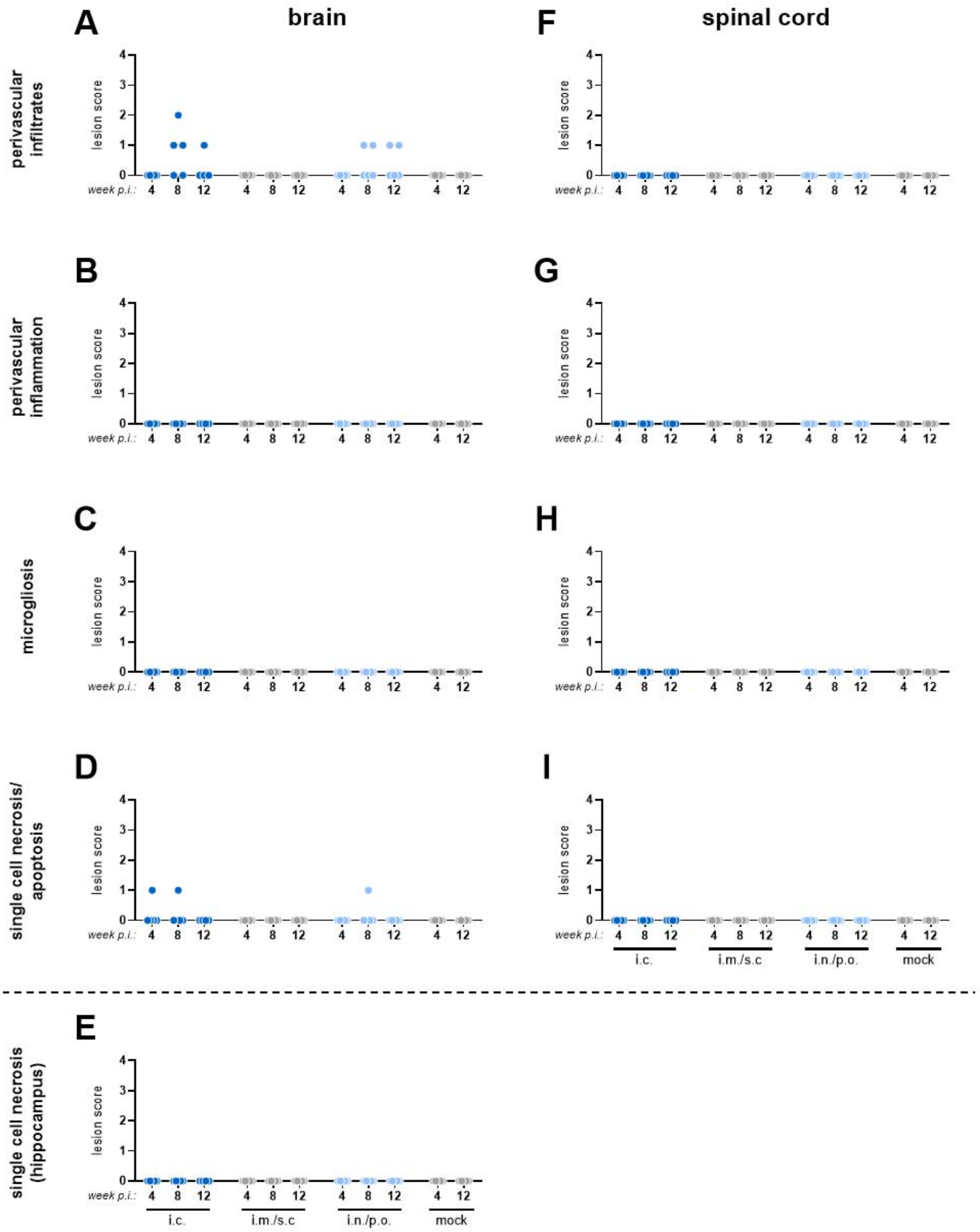


**Supplemental Figure S5. Individual microscopic lesion scores in the central nervous system of RusV-inoculated wood mice (Exp. 3).** Adult wood mice were RusV-inoculated via the indicated routes. Five animals each per group were euthanized at four, eight or twelve weeks post infection (p.i.). The presence and severity of microscopic lesions was semiquantitatively scored by blinded evaluation. Score 0: lesion not detectable, 1: minimal (<5% of the area are affected or 1-3 foci), 2: mild (6%–40% or >3 foci), 3: moderate (41%–80%, or coalescing), 4: severe (>80%, or diffuse). Results are presented as individual scores for the indicated parameters in brain and spinal cord. Coloured dots represent RusV-positive animals, while grey dots represent RusV-negative animals, as determined by RT-qPCR. i.c. = intracranial; i.m./s.c. = intramuscular/subcutaneous; i.n./p.o. = intranasal/peroral. Raw data underlying this figure are provided as Supplementary Table S5.

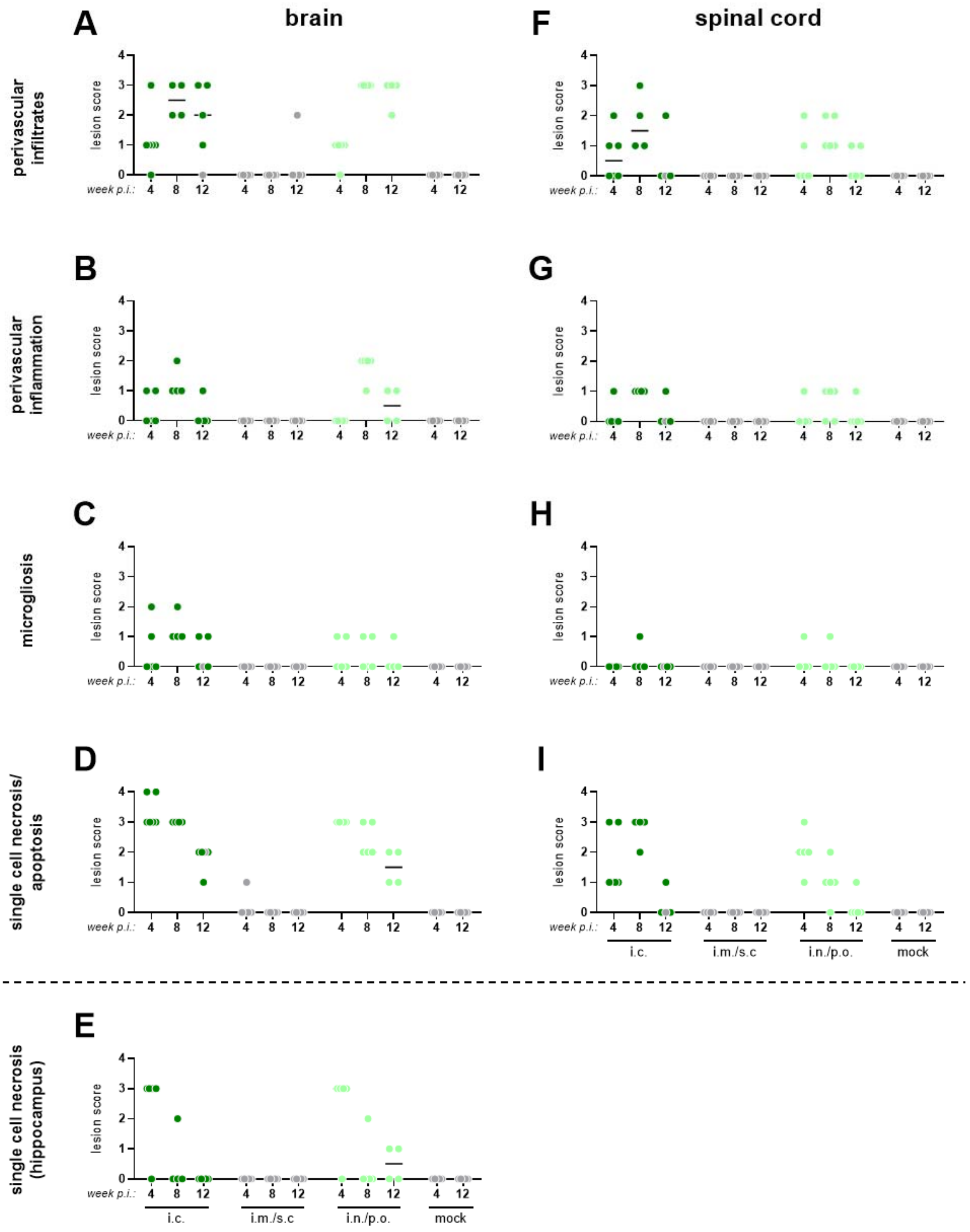


**Supplemental Figure S6. Individual microscopic lesion scores in the central nervous system of RusV-inoculated rats (Exp. 3).** Juvenile Lewis rats were RusV-inoculated via the indicated routes. Five animals each per group were euthanized at four, eight or twelve weeks post infection (p.i.). The presence and severity of microscopic lesions was semiquantitatively scored by blinded evaluation. Score 0: lesion not detectable, 1: minimal (<5% of the area are affected or 1-3 foci), 2: mild

(6%–40% or >3 foci), 3: moderate (41%–80%, or coalescing), 4: severe (>80%, or diffuse). Results are presented as individual scores for the indicated parameters in brain and spinal cord. Coloured dots represent RusV-positive animals, while grey dots represent RusV-negative animals, as determined by RT-qPCR. i.c. = intracranial; i.m./s.c. = intramuscular/subcutaneous; i.n./p.o. = intranasal/peroral. Raw data underlying this figure are provided as Supplementary Table S6.

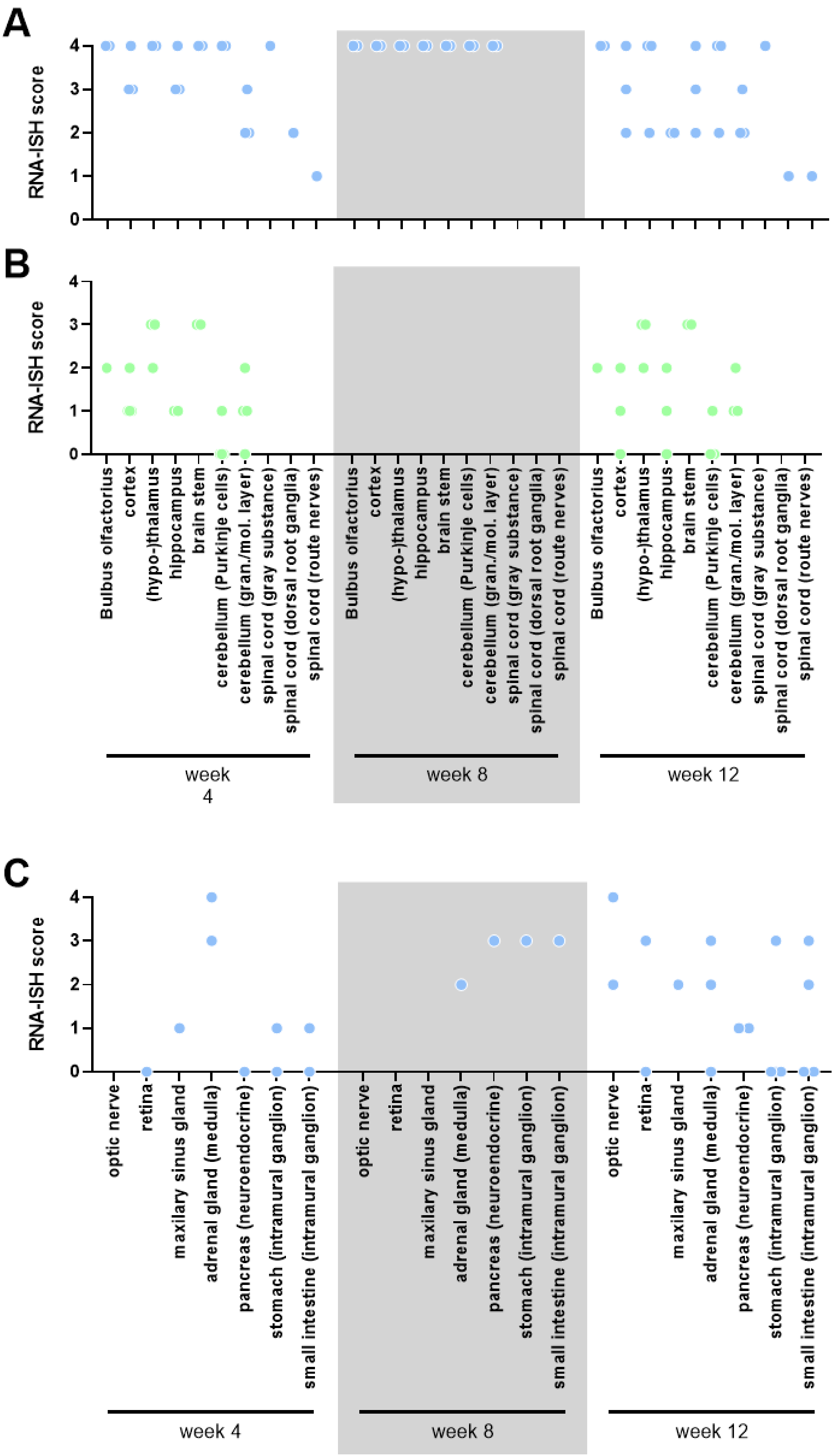
A
B
C
RNA-ISH score
0
1
2
3
4
Bulbus olfactorius
cortex
(hypo-)thalamus
hippocampus
brain stem
cerebellum (Purkinje cells)
cerebellum (gran./mol. layer)
spinal cord (gray substance)
spinal cord (dorsal root ganglia)
spinal cord (route nerves)
week 4
week 8
week 12
optic nerve
retina
maxilary sinus gland
adrenal gland (medulla)
pancreas (neuroendocrine)
stomach (intramural ganglion)
small intestine (intramural ganglion)

**Supplemental Figure S7. Detection of RusV RNA by *in situ* hybridization (RNA-ISH) in RusV-inoculated wood mice and rats (Exp. 3).** Adult wood mice and juvenile Lewis rats were inoculated via the indicated routes with a pool of RusV-positive brain homogenates from experimentally inoculated rats from groups D1 and D2 of Exp. 2. Central nervous system tissue of selected intranasally/perorally (i.n./p.o.) inoculated wood mice (A) or rats (B) and peripheral organs of selected i.n./p.o.-inoculated wood mice (C) euthanized at the indicated weeks post infection (p.i.) were analysed. The presence and extend of viral RNA is presented by semiquantitative scores. Only peripheral tissues with detection of viral RNA in at least one animal are shown. Raw data underlying this figure are provided as Supplementary Table S7 and S8.

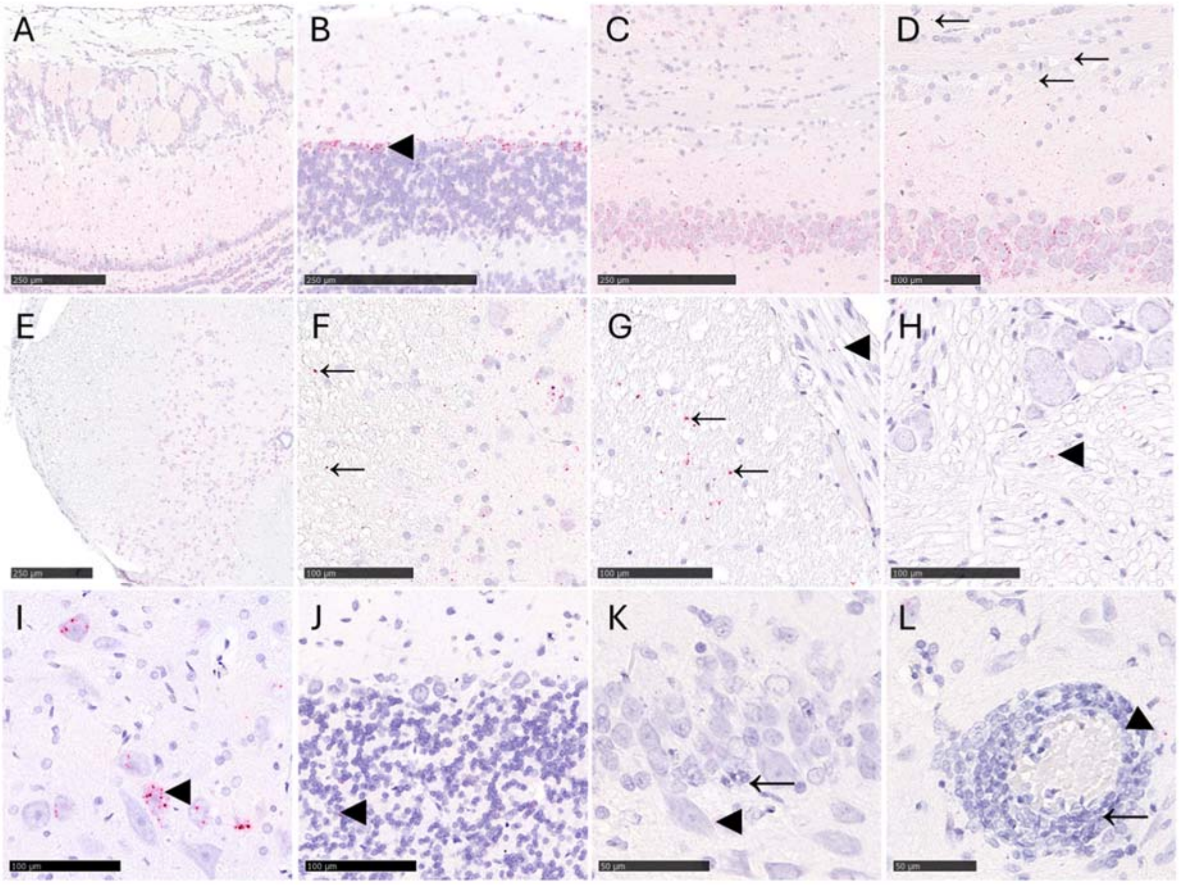


**Supplemental Figure S8. RusV RNA detection in the brain and spinal cord of wood mice (A-H) and rats (I-L) by RNA *in situ* hybridization (RNA-ISH; Exp. 3).** (A-H) Wood mice: RusV RNA in all layers of the olfactory bulb (A), cerebellum, particularly in Purkinje cells (◄, B), hippocampus (C, D), spinal cord (E, F), in dorsal root nerves (◄, G), and dorsal root ganglia (◄, H). Note the detection of RusV RNA mainly in neurons but less abundant in the white substance (←), and the absence of inflammation, while large amounts of RusV RNA are present (D, F, G, H). (I-L) Rats: Detection of abdundant RusV RNA by RNA-ISH in neurons (◄) of the cerebellar nuclei (I), less abundant in neurons (◄) of the cerebellar granular layer neurons (J), scattered in neurons of the hippocampus (◄, K) and in the neuropil (◄, L). Note that intralesional presence of some RusV RNA was only found in few locations, exemplarily shown for the hippocampus with neuronal necrosis (←, K) and adjacent to perivascular infiltrates with associated inflammation (←, L). Bar: 50, 100 or 250 µm, as indicated.

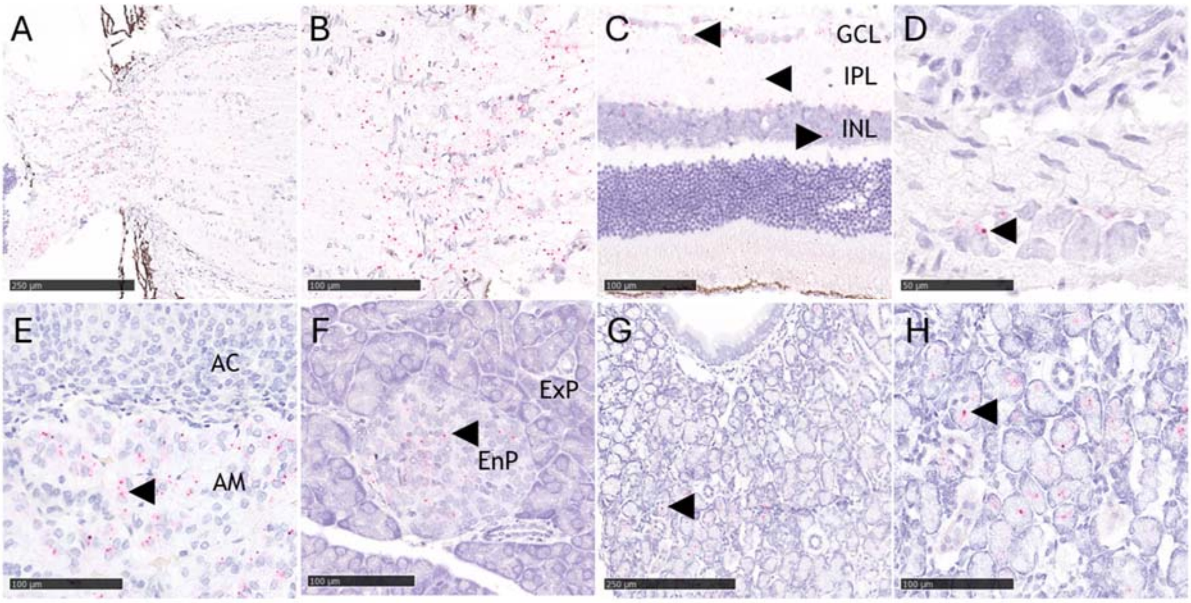


**Supplemental Figure S9. RusV RNA detection in tissues beyond brain and spinal cord of wood mice by RNA *in situ* hybridization (RNA-ISH; Exp. 3).** RusV RNA in the optic nerve (A, B), in the ganglion cell layer (GCL), inner plexiform layer (IPL), inner nuclear layer (INL) of the retina (◄, C), in intramural ganglion neurons (◄) of the intestine (D), in neuroendocrine cells (◄) of the adrenal medulla (AM) but not in epithelial cells of the adrenal cortex (AC) (E), in cells (◄) of the endocrine pancreas (EnP) but not in the exocrine pancreatic (ExP) epithelium (F), in the glandular epithelium (◄) of the maxillary sinus gland (G, H). Bar: 50, 100 or 250 µm, as indicated.
